# Inland-coastal bifurcation of southern East Asians revealed by Hmong-Mien genomic history

**DOI:** 10.1101/730903

**Authors:** Zi-Yang Xia, Shi Yan, Chuan-Chao Wang, Hong-Xiang Zheng, Fan Zhang, Yu-Chi Liu, Ge Yu, Bin-Xia Yu, Li-Li Shu, Li Jin

**Author notes:** Corresponding Authors: Shi Yan, Li Jin.

## Abstract

The early history of the Hmong-Mien language family and its speakers is elusive. A good variety of Hmong-Mien-speaking groups distribute in Central China. Here, we report 903 high-resolution Y-chromosomal, 624 full-sequencing mitochondrial, and 415 autosomal samples from 20 populations in Central China, mainly Húnán Province. We identify an autosomal component which is commonly seen in all the Hmong-Mien-speaking populations, with nearly unmixed composition in Pahng. In contrast, Hmong and Mien respectively demonstrate additional genomic affinity to Tibeto-Burman and Kra-Dai speakers. We also discover two prevalent uniparental lineages of Hmong-Mien speakers. Y-chromosomal haplogroup O2a2a1b1a1b-N5 diverged ∼2,330 years before present (BP), approximately coinciding with the estimated time of Proto-Hmong-Mien (∼2,500 BP), whereas mitochondrial haplogroup B5a1c1a significantly correlates with Pahng and Mien. All the evidence indicates a founding population substantially contributing to present-day Hmong-Mien speakers. Consistent with the two distinct routes of agricultural expansion from southern China, this Hmong-Mien founding ancestry is phylogenetically closer to the founding ancestry of Neolithic Mainland Southeast Asians and present-day isolated Austroasiatic-speaking populations than Austronesians. The spatial and temporal distribution of the southern East Asian lineage is also compatible with the scenario of out-of-southern-China farming dispersal. Thus, our finding reveals an inland-coastal genetic discrepancy related to the farming pioneers in southern China and supports an inland southern China origin of an ancestral meta-population contributing to both Hmong-Mien and Austroasiatic speakers.

## INTRODUCTION

Starting ∼9,000 years before present (BP), China is the second earliest birthplace of agriculture, following the Near East^1,2^. Subsequently, farming dispersed into adjacent areas of East Asia, especially Southeast Asia, the Korean Peninsula, and the Japanese archipelago. However, it is still not fully settled whether and to what extent human migration propelled the dispersal of agriculture. Particularly for Southeast Asia, recent archaeogenetic studies support that the East Asian ancestry of the first farmers in Southeast Asia can be traced to southern China^3,4^, raising the further issue on the deeper demographic prehistory of both regions.

The movement of farming populations has also been presumed to facilitate the expansion of their languages^5^. Seven major language families in East Asia are involved in this hypothesis: Austroasiatic, Austronesian, Hmong-Mien, Japonic, Kra-Dai (formerly known as Tai-Kadai), Koreanic, and Sino-Tibetan^6^. Compared with the others, the Hmong-Mien language family lacked writing system until the recent centuries, hence its notably obscure history, and most of the history was conserved either by oral legends, or literature documented in Chinese, which often confused different populations. Therefore, linguistic and genetic studies are crucial to the reconstruction of the Hmong-Mien prehistory. Some linguists have attempted to connect Hmong-Mien with other language families genealogically, but none of the hypotheses are widely accepted^7^. Likewise, in contrast to previous studies based on mitochondrial^8^ or Y-chromosomal^9^ data, there has been no genome-wide studies concentrating on Hmong-Mien-speaking populations up to date.

Currently, Hmong-Mien speakers mainly dwell in South Central and Southwest China, with sporadic distribution in Mainland Southeast Asia (MSEA) due to the migration within the recent centuries^7^. The linguistic phylogeny of the Hmong-Mien languages remains controversial, while a common hypothesis categorizes them into two sub-branches: Hmongic and Mienic. Whereas the Hmongic languages receive more influence from Tibeto-Burman languages, the Mienic languages show more impact from Chinese. In Húnán Province of Central China, both clades distribute, including two Hmongic languages: Hmong (to be specific, Qo Xiong) in the northwest Húnán and Pahng (Hm Nai) in the southwest, as well as a Mienic language (Mien, or specifically, Iu Mien) in the southeast. Along with a high variety of other Sino-Tibetan-and Kra-Dai-speaking groups (Fig. 1B), especially Sinitic Tǔhuà and Wǎxiāng (known for its retentions of Old Chinese^10^), Tǔjiā of Tibeto-Burman, and Kam of Kam-Sui, Central China is an appropriate field to depict a fine-scale genetic landscape of Hmong-Mien speakers with neighboring populations.

**Figure 1.**
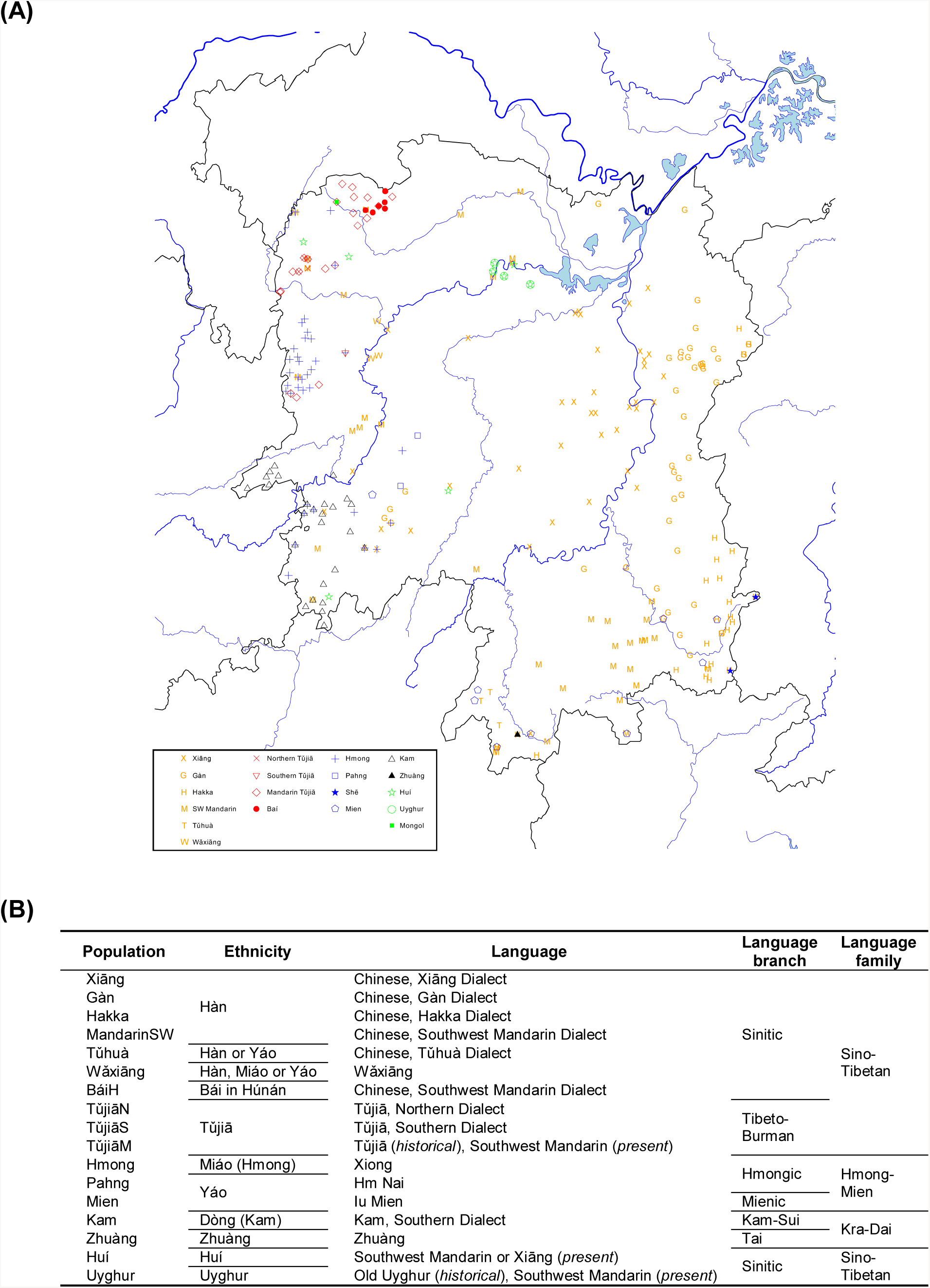
Geographic and linguistic information of sampled populations. **(A)** Geographic distribution of populations in Central China. Locations are accurate to township level. **(B)** Recognized ethnicity and linguistic affiliation of every population.

## RESULTS

To comprehensively represent the genetic diversity of Central China, we sampled and genotyped 903 high-resolution Y-chromosomal samples, 624 full-sequencing mitochondrial samples, and 415 autosomal samples from Central China, mainly in Húnán Province, covering all the 11 officially recognized ethnic groups of this region (Fig. 1). Given the criteria of both the officially recognized ethnicity and linguistic affiliation, we subdivided these groups into 20 populations. We then analyzed our data along with genetic data of ancient and modern populations from published studies (see Method).

### Genetic structure of Húnán Province

The pattern of principal component analysis (PCA) of East-Asia-wide populations^11^ (Fig. 2B) shows that at least three meta-populations contribute to the genomic diversity of populations in Húnán: one represented by northern East Asians [Northern Hàn, Mongolic and Tungusic speakers, Japanese, Korean, Tibeto-Burmans (except Lāhù and Burmese), and the Devil’s Cave sample (referred as “Devil’s Cave” for short below)^12^; right], one by southeast Asians [Austronesians, Austroasiatic and Kra-Dai speakers, and a Neolithic Laotian individual (labelled as “Laos_LN_BA”)^3^; left], and one by Hmong-Mien-speaking Hmong and Pahng (top). Compared with other Kra-Dai-(Dǎi and Thai) and Tibeto-Burman-speaking (Tibetan, Sherpa, Nàxī, and Yí) populations, Kam and Tǔjiā in Húnán shift towards Hmong and Pahng, suggesting an additional genomic impact on them from Hmong-Mien speakers, or/and gene flow between Kam and Tǔjiā. Including subpopulations of Hàn in Húnán (Gàn, Xiāng, Hakka, and Southwest Mandarin), Southern Hàn clusters between northern East Asians and Southeast Asians, consistent with the “Yellow/Yangtze River Ghost Population” theory and the inference that Hàn Chinese is the admixture of both proposed populations^13^.

**Figure 2.**
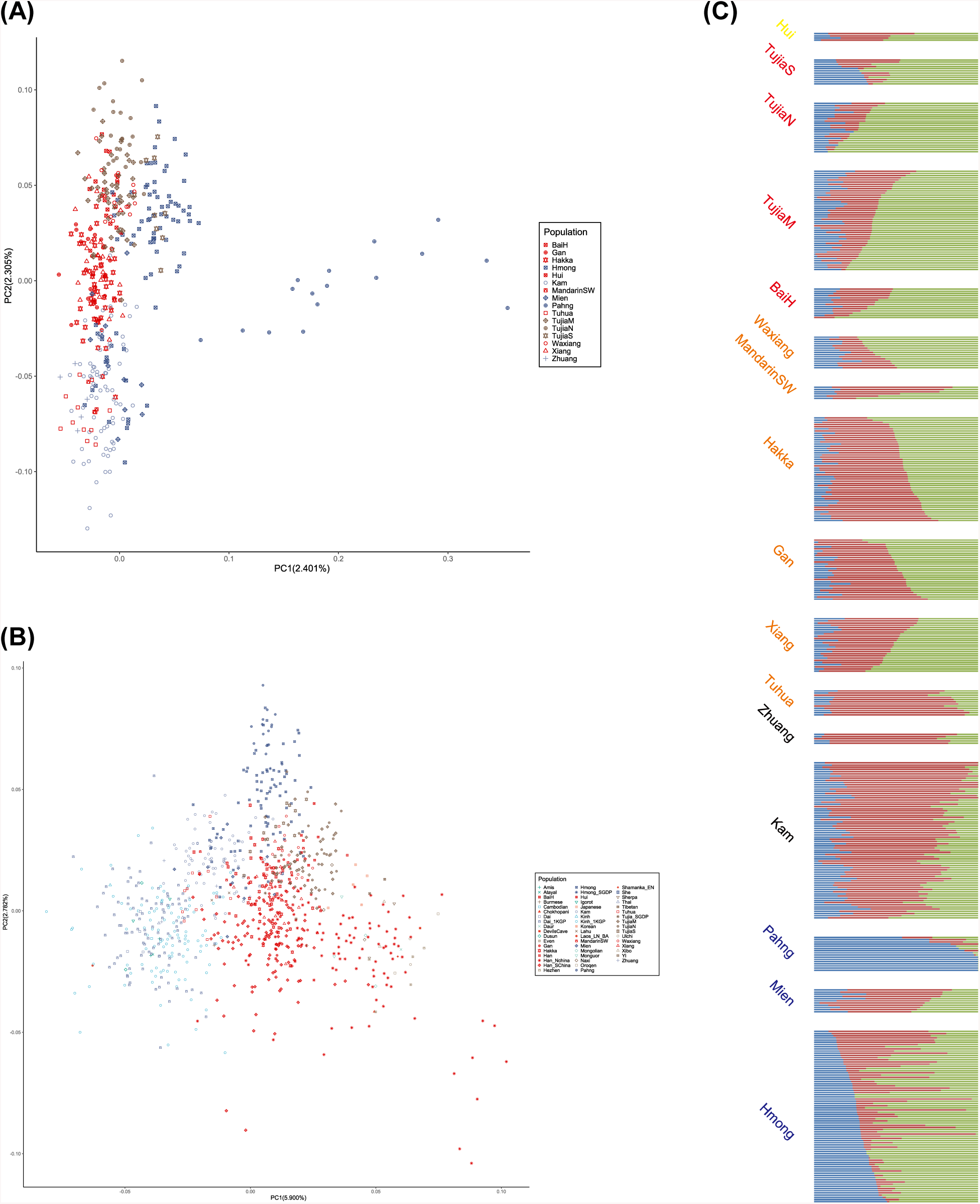
Genomic structure of Húnán in relation to East-Asia-wide diversity. **(A** to **B)** Principal component analysis (PCA) (A) restricted to populations in Húnán and (B) together with a wide-ranging panel of 473 modern East Asians and four projected ancient samples. **(C)** Model-based clustering equipped with ADMIXTURE. When K=3, there is a high intra-population consistency of genetic composition, and the distribution of ancestral components is highly correlated with linguistic affiliation: blue, Hmong-Mien; green, Sino-Tibetan; red, Kra-Dai.

To present a finer-scale genomic structure within this region, we performed PCA^11^ (Fig. 2A) and model-based clustering^14^ (K = 3, Fig. 2C) exclusively for populations in Húnán. We observe that the genomic structure is highly associated with the linguistic affiliation of populations. Northern Tǔjiā, Kam, and Pahng construct three extremes in the PCA of populations in Húnán (top, bottom, and right, respectively, Fig. 2A), matching their highest proportion of ancestral components respectively related to Sino-Tibetan, Kra-Dai, and Hmong-Mien in ADMIXTURE plot (green, red, and blue, respectively, Fig. 2C). Gàn, Xiāng, and Hakka are modeled with nearly equivalent proportion of Sino-Tibetan-and Kra-Dai-related ancestral components (Fig. 2C) and cluster with each other in the PCA plot (Fig. 2A) with non-positive pairwise F_ST_ to each other (all < 0.000, Extended Data Fig. 1), indicating that intermarriage among main subpopulations of Hàn is not significantly fewer than marriage within a subpopulation. On the contrary, Wǎxiāng and Tǔhuà significantly deviate from the main subpopulations of Hàn and cluster with Tǔjiā and Kam in PCA (Fig. 2A) and TreeMix^15^ (Fig. 3A) plot and harbor additional Sino-Tibetan-and Kra-Dai-related components in PCA plot (Fig. 2C), respectively, consistent with the particularity of their languages distinct from other Sinitic ones.

**Figure 3.**
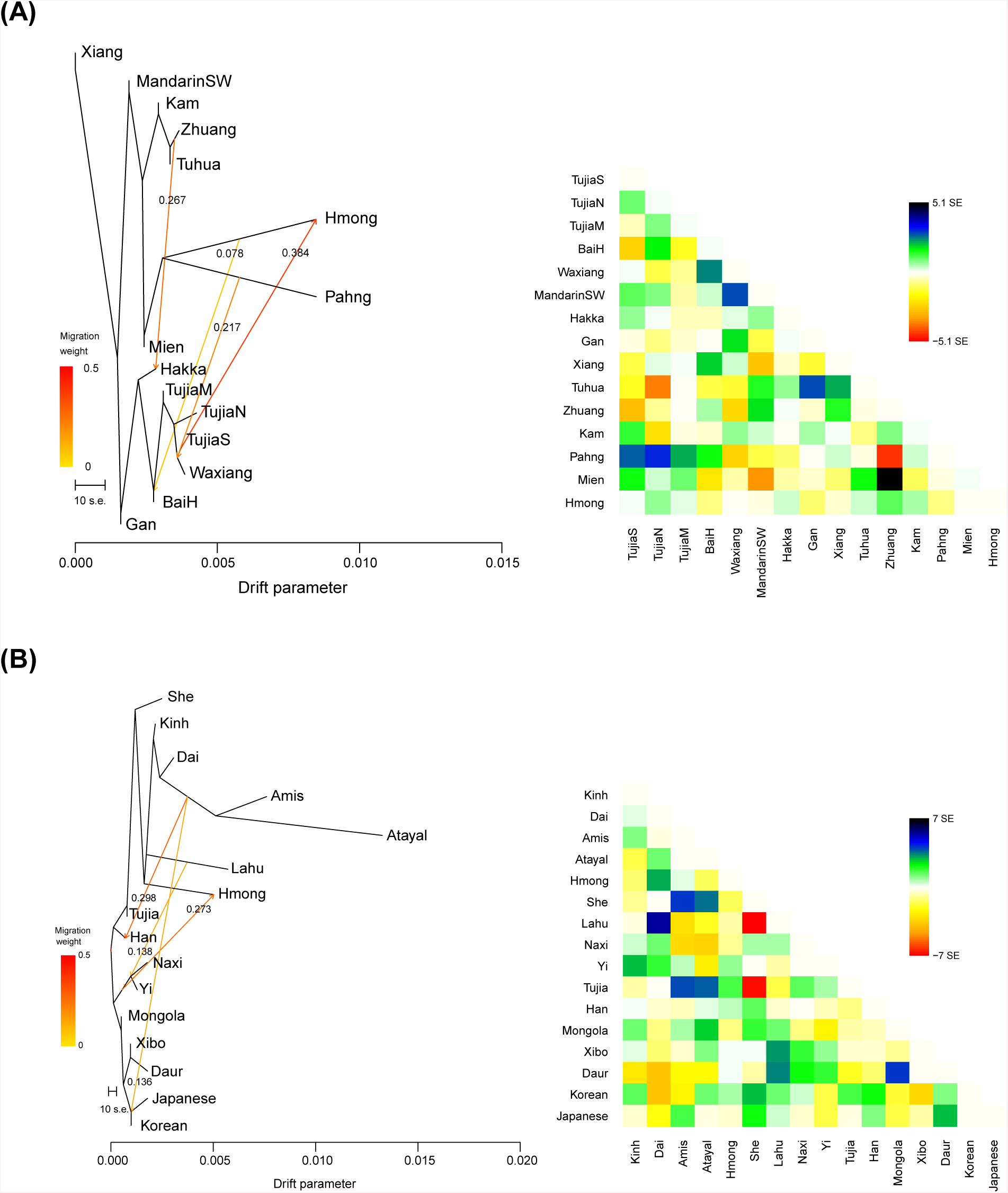
Unrooted TreeMix analysis modelling a tentative phylogenetic relationship. TreeMix graph of **(A)** populations in Húnán and **(B)** East-Asia-wide populations.

Focusing on the genomic relationship among Hmong-Mien speakers, we find that Pahng, Hmong, and Mien cluster together in the TreeMix plot for populations in Húnán (Fig. 3A). In ADMIXTURE plot (Fig. 2C), Pahng is modeled with a nearly unmixed form of Hmong-Mien-related component, consistent with the results of admixture *f*_*3*_-statistics (Table 1) and pairwise F_ST_ (Extended Data Fig. 1). The significant positive values of all the *f*_*3*_(Pahng; Hmong, Other Central Chinese)^16^ (all Z > 14.7) support that Pahng has not received substantial gene flow from non-Hmong-Mien-speaking populations and shows that Pahng has undergone a strong and unique genetic drift. Supporting the latter inference, all the pairs of F_ST_ between Pahng and other populations (all ≥ 0.018) are higher than the ones among other population themselves (all ≤ 0.017). The analyses above confirm that Pahng represents a Hmong-Mien-sharing ancestry that is absent, or at least with fewer amounts, in non-Hmong-Mien populations. Corresponding to the intermediate position between Pahng and Tǔjiā in the PCA plot (Fig. 2A) and a substantial contribution of Sino-Tibetan-related component in ADMIXTURE plot (Fig. 2C), we find an additional migration to Hmong from Tibeto-Burman source modeled by TreeMix (Nàxī and Yí, Fig. 3B), supporting an additional genomic affinity of Hmong to Tibeto-Burman speakers. Meanwhile, Mien approximately clusters with Kam in the PCA plot (Fig. 2A), confirming that Mien carries both Hmong-Mien-related and Kra-Dai-related ancestries. Other than Hmong-Mien-speaking populations, Southern Tǔjiā is the only one who harbors significant Hmong-Mien-related ancestral component (Fig. 2C), which is explainable due to its geographic proximity to Hmong and possible intermarriage (Fig 1A). Key findings of admixture *f*_*3*_-statistics of populations in Húnán (Table 1) are compatible with the inference above. Given that some Sinitic populations, especially Tǔhuà, also have a substantial Kra-Dai-related ancestry, we suggest that the dispersal and subsequent admixture associated with Hmong-Mien-related ancestry are highly parallel to the dissemination and language contact of different Hmong-Mien languages.

**Table 1.** Admixture *f*_*3*_ statistics of Central China populations. The key discoveries revealed by corresponding *f*_*3*_ statistics are mentioned in the header. *f*_*3*_ statistics with a relatively negative value (< −4.0×10^-5^) and Z-score (Z < −0.5, one-tailed *p* < 0.309) are shaded.

| Populations | Pahng does not have significant non-Hmong-Mien ancestry |  | Hmong has extra Sino-Tibetan-related ancestry |  | Mien has both Kra-Dai- and Hmong-Mien-related ancestries |  |  |  | TujiaS has extra Hmong-Mien-related ancestry |  | Han in Hunan has both Sino-Tibetan- and Kra-Dai-related ancestries |  |  |  |  |  |
| --- | --- | --- | --- | --- | --- | --- | --- | --- | --- | --- | --- | --- | --- | --- | --- | --- |
| | $f_3(\text{Pahng; Hmong, X})$ | | $f_3(\text{Hmong; Pahng, X})$ | | $f_3(\text{Mien; Pahng, X})$ | | $f_3(\text{Mien; Hmong, X})$ | | $f_3(\text{TujiaS; TujiaN, X})$ | | $f_3(\text{Xiang; TujiaN, X})$ | | $f_3(\text{Gan; TujiaN, X})$ | | $f_3(\text{Hakka; TujiaN, X})$ | |
|  | mean | Z-score | mean | Z-score | mean | Z-score | mean | Z-score | mean | Z-score | mean | Z-score | mean | Z-score | mean | Z-score |
| Xiang | 0.01854 | 16.911 | 0.00125 | 2.109 | 0.00035 | 0.401 | 0.00075 | 1.015 | 0.00186 | 1.876 | n/a | n/a | -0.00060 | -0.875 | -0.00041 | -0.689 |
| Gan | 0.01825 | 17.307 | 0.00128 | 2.094 | 0.00044 | 0.480 | 0.00071 | 0.957 | 0.00117 | 1.180 | -0.00124 | -2.264 | n/a | n/a | -0.00154 | -2.636 |
| Hakka | 0.01906 | 18.564 | -0.00002 | -0.043 | -0.00017 | -0.171 | 0.00062 | 0.746 | 0.00144 | 1.539 | 0.00046 | 0.728 | -0.00022 | -0.324 | n/a | n/a |
| MandarinSW | 0.01949 | 16.510 | 0.00104 | 1.544 | 0.00093 | 0.876 | 0.00179 | 2.202 | 0.00096 | 0.911 | 0.00053 | 0.827 | -0.00016 | -0.254 | -0.00036 | -0.546 |
| Waxiang | 0.02084 | 16.973 | -0.00044 | -0.759 | 0.00106 | 1.060 | 0.00343 | 3.930 | 0.00092 | 1.051 | 0.00342 | 4.520 | 0.00279 | 3.246 | 0.00224 | 3.571 |
| Tuhua | 0.01814 | 16.581 | 0.00176 | 3.265 | -0.00026 | -0.249 | -0.00052 | -0.633 | 0.00019 | 0.163 | -0.00296 | -4.489 | -0.00270 | -3.499 | -0.00285 | -3.909 |
| BaiH | 0.01957 | 16.032 | -0.00002 | -0.037 | 0.00159 | 1.554 | 0.00293 | 3.392 | 0.00343 | 3.530 | 0.00207 | 2.743 | 0.00297 | 4.047 | 0.00187 | 3.336 |
| TujiaM | 0.01987 | 17.592 | -0.00045 | -0.805 | 0.00023 | 0.258 | 0.00159 | 1.957 | 0.00264 | 3.045 | 0.00315 | 3.468 | 0.00326 | 3.732 | 0.00230 | 4.393 |
| TujiaN | 0.01991 | 17.382 | -0.00043 | -0.606 | 0.00066 | 0.663 | 0.00222 | 2.675 | n/a | n/a | n/a | n/a | n/a | n/a | n/a | n/a |
| TujiaS | 0.02031 | 17.996 | 0.00027 | 0.454 | 0.00263 | 2.510 | 0.00375 | 4.121 | n/a | n/a | 0.00387 | 4.047 | 0.00418 | 4.559 | 0.00315 | 5.160 |
| Hmong | n/a | n/a | n/a | n/a | 0.00132 | 1.505 | n/a | n/a | -0.00002 | -0.019 | 0.00082 | 1.334 | 0.00161 | 2.280 | 0.00066 | 1.374 |
| Pahng | n/a | n/a | n/a | n/a | n/a | n/a | 0.00132 | 1.505 | -0.00094 | -0.863 | -0.00030 | -0.426 | 0.00007 | 0.083 | 0.00012 | 0.164 |
| Mien | 0.01813 | 15.537 | 0.00172 | 2.735 | n/a | n/a | n/a | n/a | 0.00077 | 0.685 | -0.00047 | -0.711 | -0.00032 | -0.436 | -0.00051 | -0.715 |
| Kam | 0.01726 | 16.128 | 0.00187 | 3.255 | 0.00041 | 0.444 | -0.00024 | -0.291 | -0.00038 | -0.368 | -0.00242 | -4.433 | -0.00220 | -2.996 | -0.00272 | -4.304 |
| Zhuang | 0.02046 | 14.722 | -0.00013 | -0.157 | -0.00243 | -1.932 | -0.00084 | -0.846 | 0.00177 | 1.362 | -0.00189 | -2.360 | -0.00040 | -0.456 | -0.00256 | -3.405 |

We then identify a series of motif Y-chromosomal lineages reflecting the demographic history (Fig. 4A and Extended Data Fig. 2A). Fitting their common linguistic affiliations, Hmong and Pahng share the patrilineage O2a2a1b1a1b-N5 (included in O2a2a1b-M7) with the second-highest (18/95 = 18.9%) and the highest frequency (7/21 = 33.3%), respectively. This patrilineage is also present in Shē^17^, a population in southeastern China who once spoke Hmong-Mien. The highest frequency patrilineage for Hmong, O1b1a1a2a-F5506 (23/95 = 24.2%, included in O1b1a-PK4), is also exclusive for Hmong. Three dominant patrilineages for Hàn Chinese^18^, Oα-F5, Oβ-F46, and Oγ-F11 (respectively included in O2a2b1a1-M117, O2a2b1*-M134(xM117), and O2a1b-IMS-JST002611), have a considerable proportion (57/463 = 12.3%, 44/463 = 9.5%, 58/463 = 12.5%, respectively) within main subpopulations of Hàn in Húnán, partly supporting a demic diffusion of Sinitic languages in this region. Compared with that, Wǎxiāng and Tǔhuà respectively have a noticeable frequency of Y-haplogroup N2a1-F1833 (6/39 = 15.4%, included in N-M231) and D1a2-F1531 (3/17 = 17.6%, included in D1a-M15), following their special linguistic and autosomal history. Intriguingly, Tǔjiā and Kam share Y-haplogroup Cα2a1-F5477 (included in C2a-F1067) with a high frequency (24/90 = 26.7% and 13/71 = 18.3%, respectively), despite their different linguistic affiliations, supporting a history of gene flow between both on the paternal side. In contrast, mitochondrial haplogroups present a uniform distribution among populations (Fig. 4B and Extended Data Fig. 2B), indicating the practice of exogamy on the maternal side. The very exceptions are Pahng and Mien, both Hmong-Mien-speaking, with the significant correlation with mitochondrial haplogroup B5 (*χ*^*2*^ = 1.03×10^-11^ and 1.73×10^-5^, respectively, whereas Bonferroni corrected *p* = 2.3×10^-4^).

**Figure 4.**
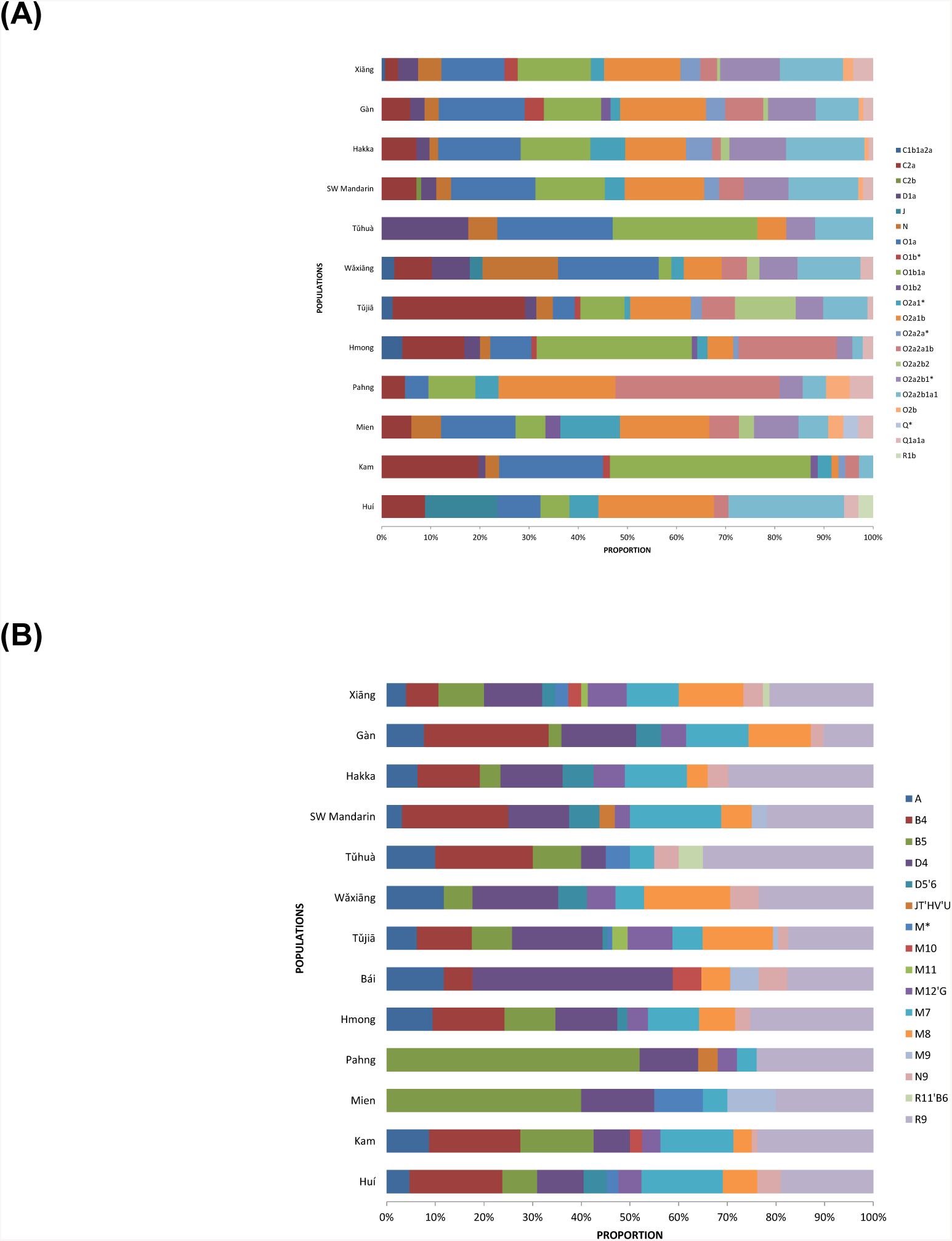
Distribution of uniparentally inherited lineages in Húnán. Proportion of **(A)** Y-chromosomal and **(B)** mitochondrial haplogroups in Central China populations.

We also performed coalescent time estimation (see Method) to calibrate the separation of some uniparentally inherited lineages (Fig. 5). Approximately synchronous with the time estimation of Proto-Hmong-Mien (∼2,500 BP)^19^, we estimate the most recent common ancestor (TMRCA) of O2a2a1b1a1b-N5 as ∼2,330 BP (Fig. 5B), consistent with slight precedence of the divergence of patrilineage than the dispersal of language and ancestral population. By phylogenetic analysis (Fig. 5A), most of the Pahng (with one exception) and Mien (with two exceptions) samples affiliated to mitochondrial haplogroup B5 are identified as the subclade B5a1c1a (TMRCA in ∼9,800 BP). Given the substantially higher frequency of Hmong-Mien speakers (63.6%, 21 out of 33) than the others in B5a1c1a, we propose a Hmong-Mien origin of this subclade.

**Figure 5.**
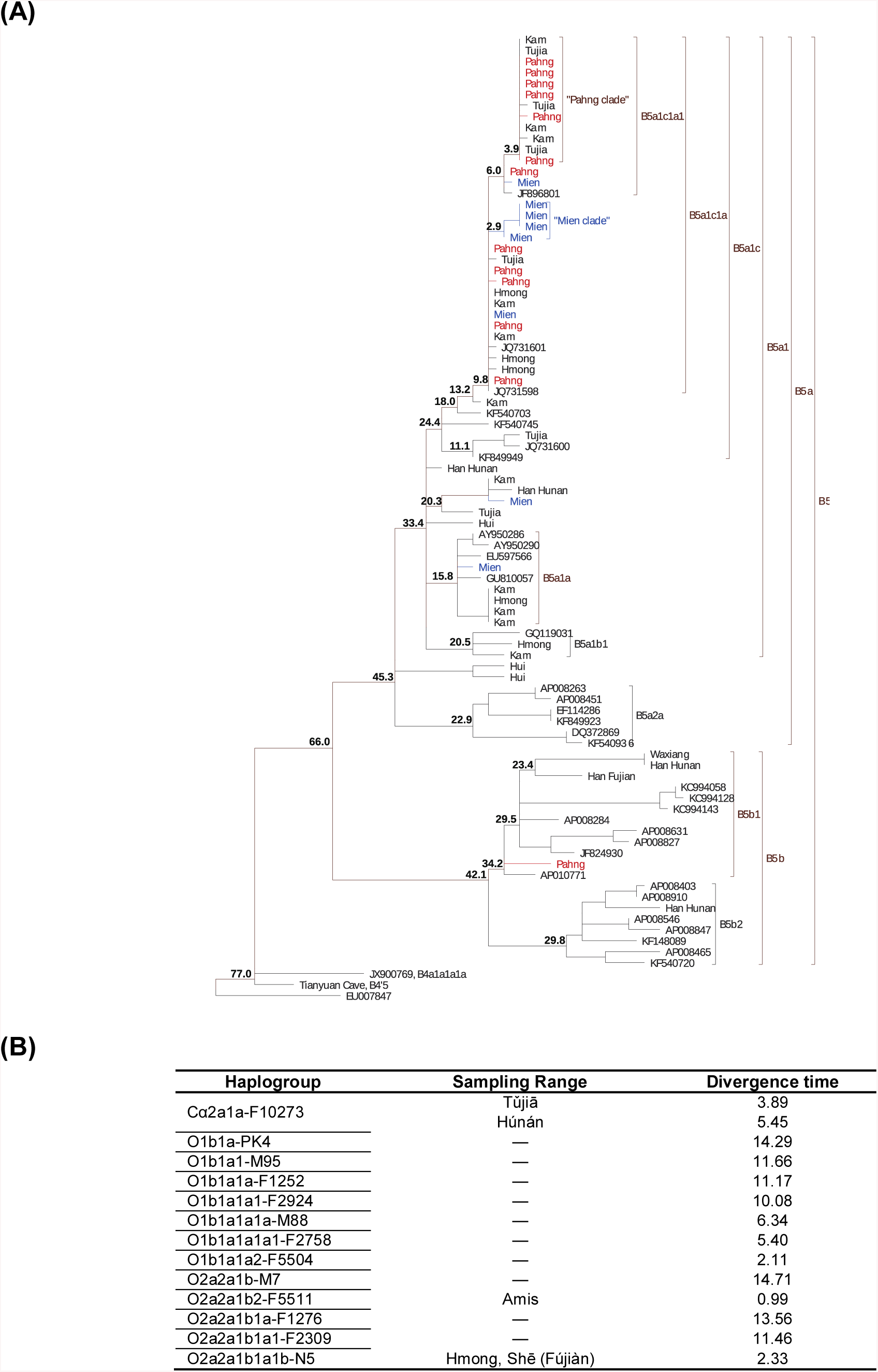
Coalescent time estimation of uniparental lineages. **(A)** Phylogeny and divergence time estimates for mitochondrial haplogroup B5. We use maximum parsimony to establish the phylogeny and Bayesian inference to estimate the divergence time (kya). **(B)** Divergence time estimates of motif Y-haplogroups.

In summary, we find strong evidence from genomic and uniparental analyses for a founding population who is responsible for the dispersal of Proto-Hmong-Mien and thus has a substantial genetic impact on present-day Hmong-Mien speakers, hence naming it as “Ancestral Hmong-Mien” (AHM).

### Inland-coastal bifurcation of southern East Asians

One of the core issues regarding the history of the Hmong-Mien language family is its place of origin. Whereas linguistic evidence, such as the reconstructed vocabulary related to wet-rice cultivation, supports a Yangtze Basin origin of Hmong-Mien^7^, archaeological record reveals a quite more complex scenario. During the Neolithic, there were two independent agricultural centers in southern China: Yangtze Delta and the middle Yangtze^1,20^ (here we definite the northern boundary of “southern China” as Qín Mountains and Huái River). The former partly explains the coastal route for farming expansion as far as Taiwan and Luzon, while the latter accounts for another inland expansion of agriculture into Southwest China and MSEA^21^. In particular, there is a strong bond between the material culture of Neolithic Yangtze Delta and later Austronesian Pacific islands previously described^22^. Nevertheless, it is still ambiguous about the relationship between Proto-Hmong-Mien and these agricultural centers. Therefore, to determine the genomic relationship between AHM and other populations is pivotal to the resolution of the issue.

Besides AHM, previous genetic studies have already distinguished two other ancestral populations highly associated with the farming dispersal from southern China^3,4,23^. One of them is represented by the East Asian ancestry of Neolithic farmers in MSEA, as well as present-day isolated Austroasiatic-speaking groups (*e.g.*, Htin and Mlabri), who largely maintain genetic continuity with the former^4^, hence responsible for Austroasiatic expansion and named as “Ancestral Austroasiatic” (AAA). The other is represented by Austronesian-speakers in Taiwan (*e.g.*, Amis and Atayal) in nearly unmixed form^23^ and accounts for the dissemination of Austronesian languages, hence addressed as “Ancestral Austronesian” (AAN).

Given that, we computed a series of *D*-statistics^16^ to address the genomic relationship between AHM and other populations (Table 2, Extended Data Table 2 & 3). Characterized by significant negative *D* (X, Mbuti; Devil’s Cave, Amis) [X = Hàn, Hmong (representing AHM), Htin/Mlabri (representing AAA); Z < −3.3 (transversion only); Extended Data Table 3], the major genomic component of Hàn, AHM, and AAA are phylogenetically closer to AAN (represented by Amis) than Devil’s Cave, hence constituting a monophyletic lineage of southern East Asians. Accordingly, we name this genomic component of Hàn as “southern East Asian ancestry of Hàn” (SEAH). Thus, Devil’s Cave is supposed to symmetrically relates to AHM, AAA, AAN, and SEAH. In view of that, Hmong harbors increasing genomic affinity to most of the northern East Asians best represented by Tibetan in addition to AHM [*D* (Tibetan, Mbuti; Hmong, Amis) > 0, Z = 7.0, Table 2], and Hàn receives slightly more genomic influence from northern East Asians than Hmong [*D* (X, Mbuti; Hmong, Hàn) < 0, Z = −2.9 for Nganasan, −2.7 for Devil’s Cave]. Devil’s Cave inclines towards Austronesian in Taiwan than isolated Austroasiatics in *D*-statistics [*D* (Devil’s Cave, Mbuti; Htin/Mlabri, Amis/Atayal) < 0, Z < −7.9], compatible with a deep ancestry from Australasian lineage harbored by isolated Austroasiatics as previously proposed^4^.

**Table 2.**
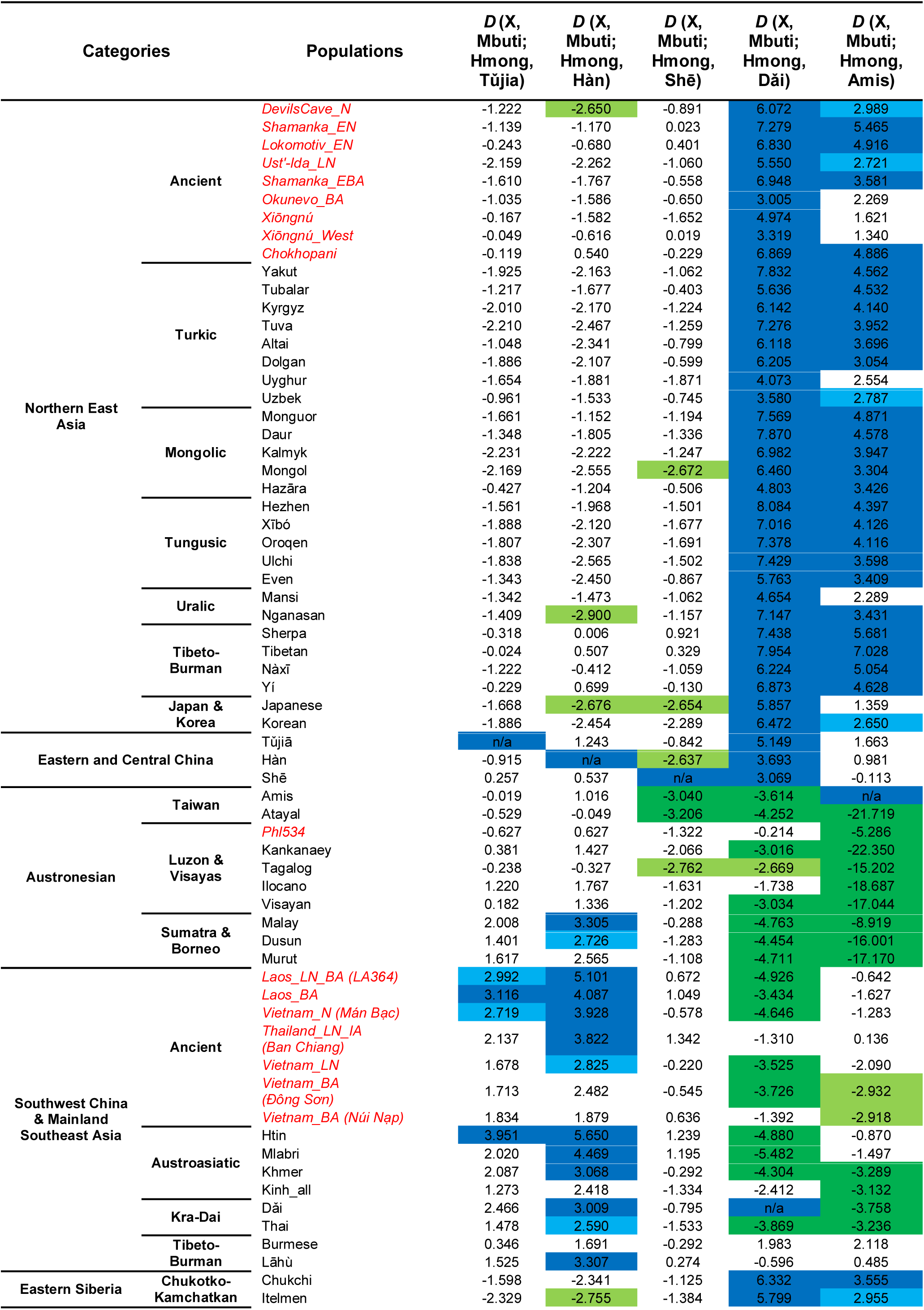
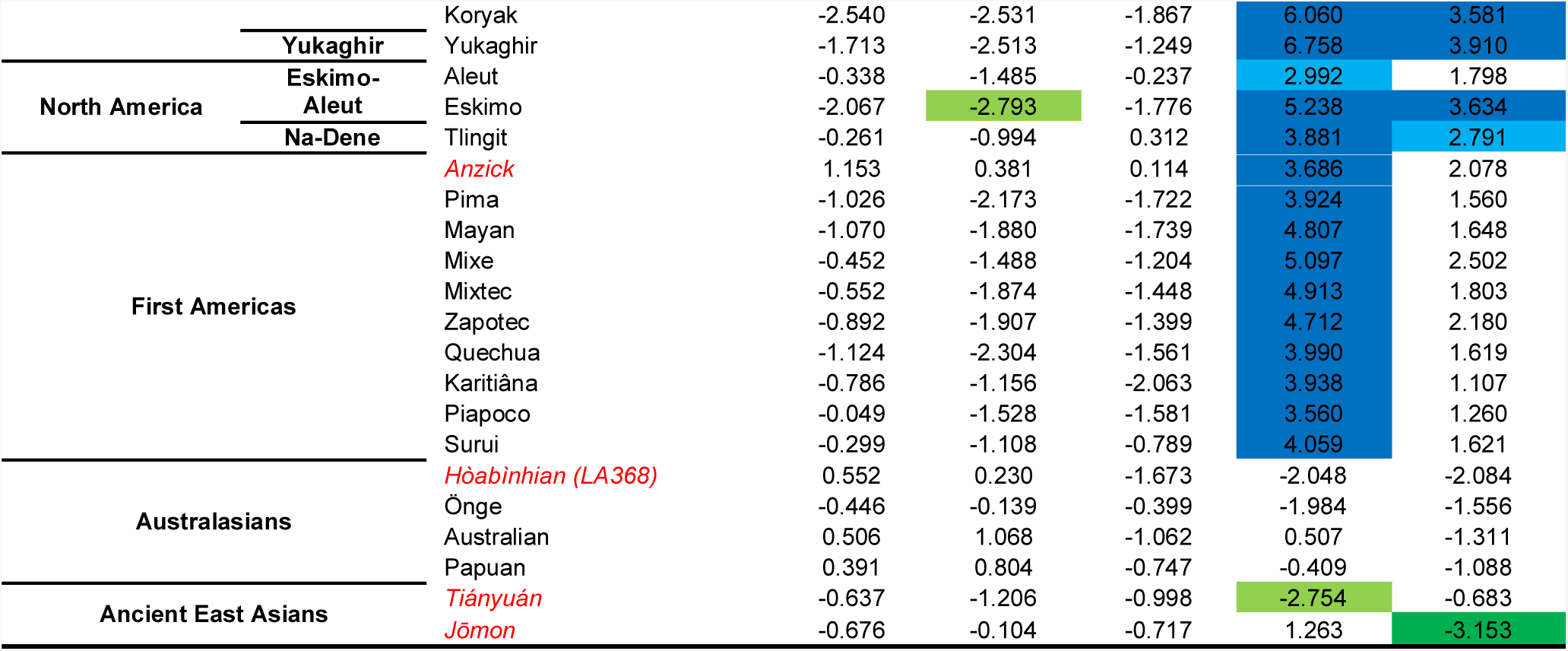
*D*-statistics detecting the major gene flow events in relation to Hmong. Significant (|Z| > 3, *i.e.*two-tailed *p* < 0.0027) and quasi-significant (2.58 < |Z| ≤ 3, *i.e.*two-tailed 0.0027 ≤ *p* < 0.01) *D*-statistics are shaded in different shade.

Revealed by the direction of *D*-statistics, different modern and ancient populations in Southeast Asia do not equally relate to Hmong, Hàn, and Amis, and can primarily categorize into three groups (Table 2). Most of Neolithic and Bronze Age samples in MSEA and isolated Austroasiatics incline towards Hmong than Hàn but symmetrically relate to Hmong and Amis. Austronesians in Luzon and Visayas highly deviate to Amis from Hmong, but symmetrically relate to Hmong and Hàn. Present-day Kra-Dai speakers and Khmer possess increase affinity both to Hmong than Hàn and to Amis than Hmong. The results above imply an intricate internal structure within the southern East Asian lineage.

Addressing that, we modeled the admixture graph^16^ to determine the phylogenetic structure of AHM, AAA, and AAN (Fig. 6). Consequently, we find the best fitting is that AAN is an outgroup to the clade constructed by AHM and AAA (Fig. 6A), supported by the lowest and feasible worst-fitting Z-score (|Z| = 2.43), the lowest final score of log-likelihood, and the positive shared drift exclusively for AHM and AAA (*f*_*2*_ = 0.006). Conversely, the other two possible cases (AHM or AAA as an outgroup, Fig. 6B and C, respectively) introduce unfitting Z-scores (|Z| = 4.54 and 4.53, respectively) and zero-length branches, indicating unfeasible assumptions about the topology. Given the current and traceable historical distribution of relevant populations, we provisionally call the lineage constructed by AHM and AAA as an “inland lineage” and the one represented by AAN as a “coastal lineage”.

**Figure 6.**
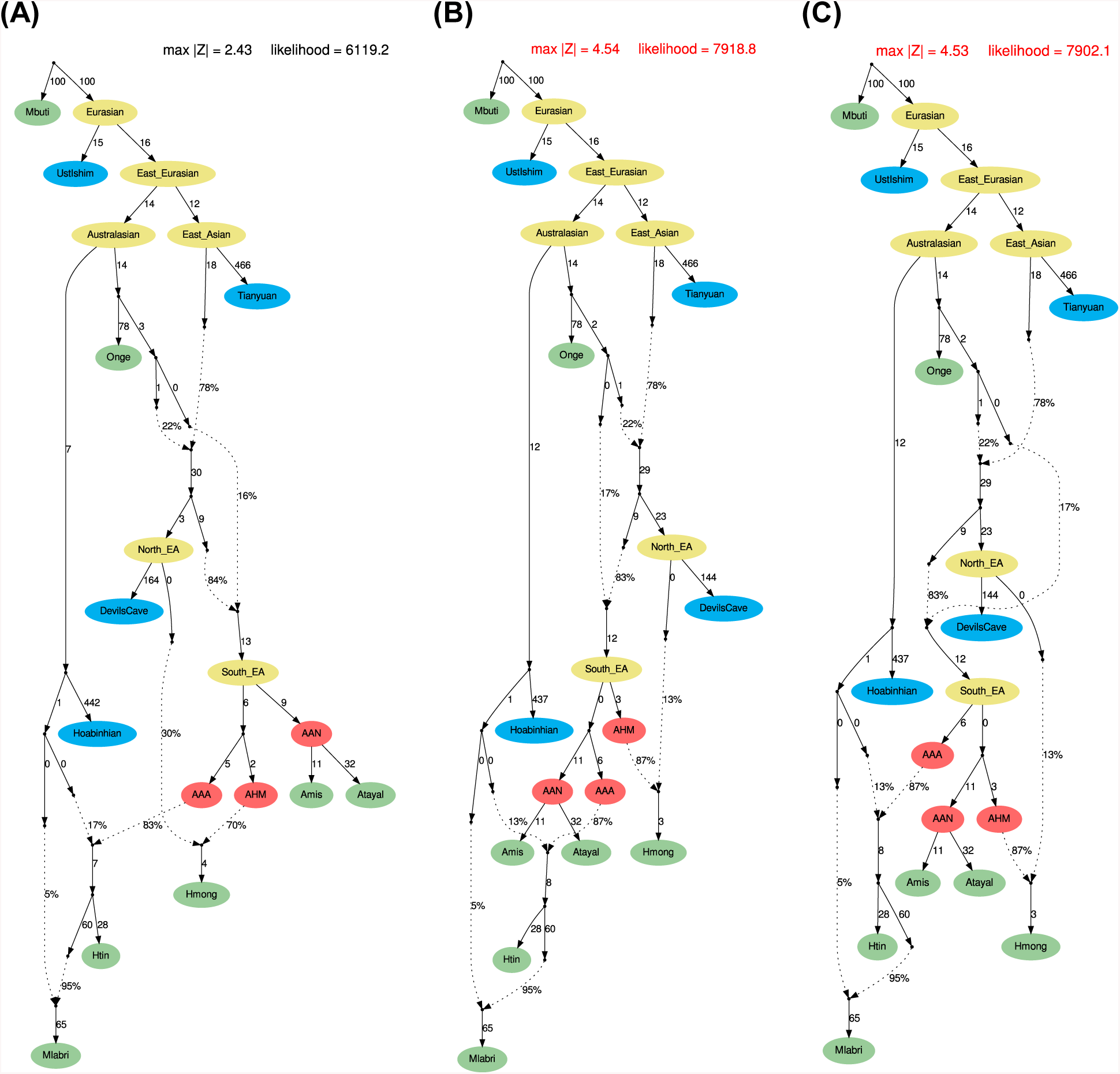
Admixture Graph modelling the phylogenetic relationship among AHM, AAA, and AAN. **(A)** AAN as the outgroup; **(B)** AHM as the outgroup; **(C)** AAA as the outgroup. Graph (A) is the best model given following criteria: (1) a feasible worst-fitting Z-score (|Z| < 3), (2) the smallest score of likelihood, and (3) without zero-length internal branch. Therefore, modeled by *qpGraph*, AHM is phylogenetically closer to AAA than AAN. We note that the worst-fitting Z-score increases [Z = 3.02 for *f*_*4*_(Tianyuan, Amis; Mlabri, Hmong)] if we do not model an increasing contribution from the Hòabìnhian lineage to Mlabri than Htin for graph (A).

To extensively model the genomic relationship of three distinct groups of Southeast Asians to the two lineages, we formally leverage the result of *D*-statistics [Supplementary Information (SI) Note 1]. Consistent with the result of admixture graph, our analysis indicates that AAA is closer to AHM than AAN (SI Note1.2). Our analysis also suggests that SEAH collectively forms a clade with AAN in contrast to AHM (SI Note 1.3) and that Second_MSEA has an increasing affinity to AAN than First_MSEA (SI Note 1.4). Considering the arrival of such an AAN-like lineage in MSEA, we find that the signal first occurred in the Bronze Age Vietnamese (dated to ∼2,000 BP) who shows an increasing affinity to Amis and a decreasing affinity to Hmong (Table 3). Given the Austro-Tai hypothesis^24^ that proposes a genealogical relationship between Kra-Dai and Austronesian languages, the occurrence of the coastal lineage in MSEA is appropriate to be explained as a southwestward migration wave related to Kra-Dai expansion^3^ after the presence of AAA in this region. Distinct from other Austronesians, both Malay and Dusun show an increasing affinity to Hmong than Hàn (Table 2). Given the closer relationship between AHM and AAA than AAN, this is also consistent with the genetic^23^, linguistic^25^, and archaeological^26^ evidence of an early Austroasiatic presence in Sumatra and Borneo. By contrast, the genomic affinity of SEAH to AAN is consistent with the language contact between Chinese and Kra-Dai languages, especially the Tai sub-branch^27^. All the evidence above indicates an earlier occurrence of the inland lineage than the coastal lineage in inland Southwest China and MSEA and supports an inland-coastal bifurcation of the southern East Asians, corresponding to the two distinct routes for agricultural expansion from southern China.

Apart from the autosomal evidence, populations sharing the inland lineage also closely connect on multiple levels. The shared vocabulary of Hmong-Mien with Austroasiatics seems to be more fundamental than one with other neighboring language families^7,28^, implying an arguably common history in an earlier stage. Regarding uniparental lineages, Hmong-Mien speakers and Austroasiatics (or Neolithic farmers in MSEA) also share a series of Y-chromosomal and mitochondrial motif haplogroups, such as Y-haplogroup O1b1a1-M95 [O1b1a1a2a-F5506 for Hmong; O1b1a1b-F789 for Malaysia_LN, Laos_LN_BA^3^, and various present-day Austroasiatic-speaking populations^29^] and O2a2a1b1-M209 [O2a2a1b1a1b-N5 for Hmong-Mien speakers; O2a2a1b1a1*-F2309(xM113, xN5) for Kinh^17^; compared with O2a2a1b2-F5511 for Amis^17^], and mitochondrial haplogroup B5a1 [B5a1c1a for Hmong-Mien speakers; B5a1a for Vietnam_N and Thailand_LN_IA^4^; B5a1c for Thailand_LN_IA^4^]. Especially for O2a2a1b-M7 (Fig. 5B), the divergence time of O2a2a1b-M7 [∼14,710 BP, TMRCA for O2a2a1b1a1-F2309 (lineage shared by Hmong-Mien and Austroasiatic speakers) and O2a2a1b2-F5511 (Amis-related lineage)] is significantly older than the divergence time of O2a2a1b1a1-F2309 (11,460 BP), consistent with the closer relationship of AHM and AAA than AAN. Besides, the divergence time for O2a2a1b1a1-F2309 and O1b1a1-M95 (∼11,660 BP) and rapid diversification [O1b1a1a-F1252, ∼11,170 BP; O1b1a1a1-F2924, ∼10,080 BP] for subgroups of the latter (Extended Data Fig. 4) may approximately synchronize with the dawn of Neolithic revolution in inland southern China (∼9,000 BP^2,21^).

### Diffusion of the southern East Asian lineage

Neolithic agriculture in Yangtze Basin is famous for its earliest domestication of sinica/japonica rice (*Oryza japonica*), which later spread into the vast tract of land ranging from Japan to Madagascar^1,30^. To test whether the agricultural diffusion from southern China is concomitant with the emigration from the same region, we used the *qpAdm*^31^ to model the spatial and temporal distribution of the southern East Asian lineage, comprising both the inland (*i.e*., AHM and AAA) and coastal (*i.e.*, AAN) lineages (Fig. 7 & Extended Data Table 4).

**Figure 7.**
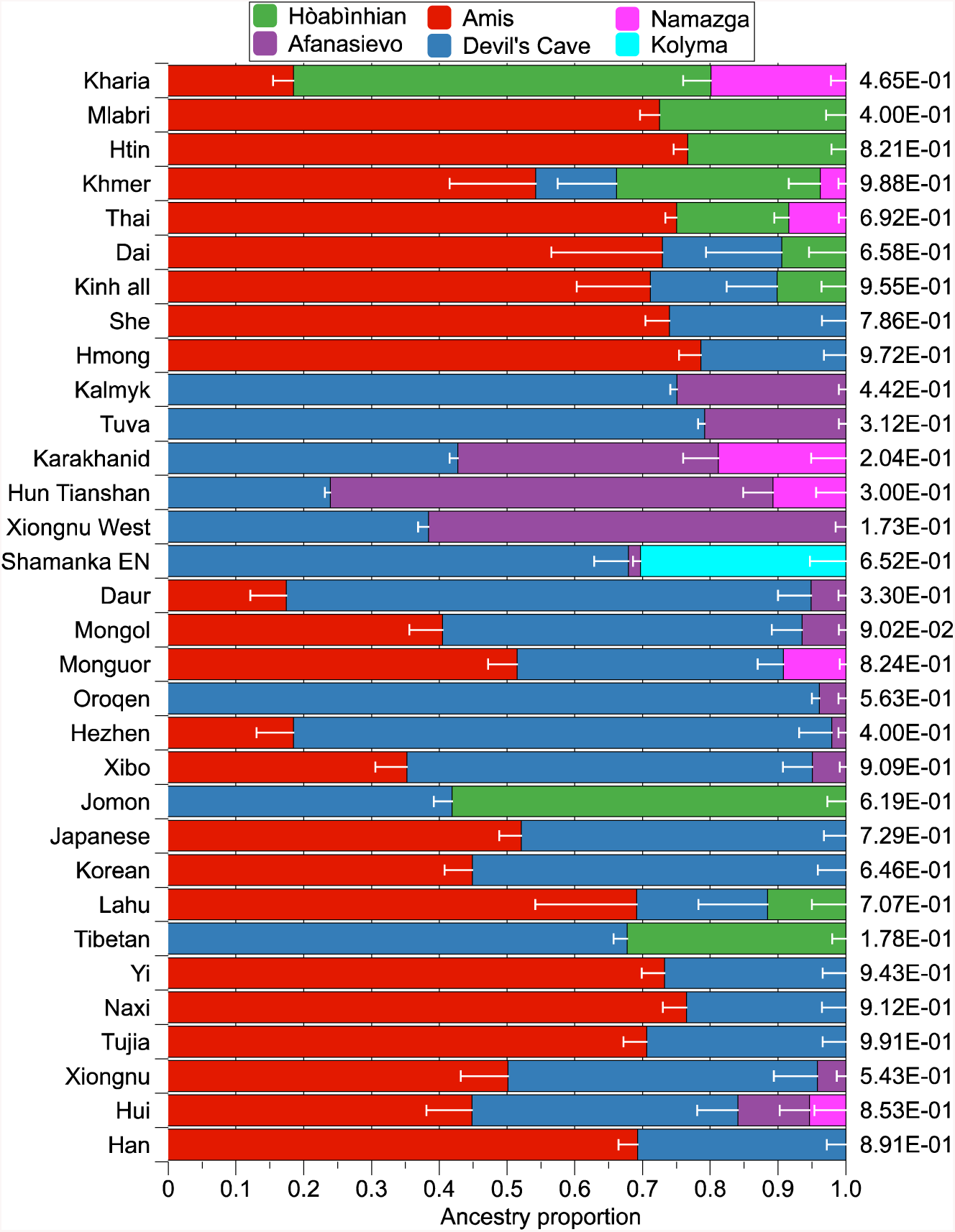
Spatial and temporal distribution of ancestries in East Asians. Reference populations and corresponding hypothesized ancestral populations: (1) Devil’s Cave (∼7,600 BP), the northern East Asian lineage; (2) Amis, the southern East Asian lineage (= AHM + AAA + AAN); (3) Hòabìnhian (∼7,900 BP), a lineage related to Andamanese and indigenous hunter-gatherer of MSEA; (4) Kolyma (∼9,800 BP), “Ancient Palaeo-Siberians”; (5) Afanasievo (∼4,800 BP), steppe ancestry; (6) Namazga (∼5,200 BP), the lineage of Chalcolithic Central Asian. Here, we report the best-fitting results of *qpAdm* based on following criteria: (1) a feasible *p*-value (> 0.05), (2) feasible proportions of all the ancestral components (mean > 0 and standard error < mean), and (3) with the highest *p*-value if meeting previous conditions.

We started from the two-way model of Amis (representing the southern East Asian lineage) and Devil’s Cave (representing the northern East Asian lineage), adding additional reference populations (Hòabìnhian, Afanasievo, Namazga, and Kolyma) when the initial model fails (see Method). In addition to MSEA populations (9.5–30.0%)^3,4^, South Asia^32,33^ (Kharia, 61.6 ± 2.2%), and the Jōmon individual (58.1 ± 2.7%) dated to ∼2,600 BP^3^, Hòabìnhian-like component also occurs in present-day Tibetans (32.3 ± 2.0%), suggesting a once wide-ranging distribution of the Australasian lineage in these regions prior to the agricultural expansion in East Asia. Regarding the East Asian lineage, there is no significant difference in fitness of Jōmon using either Devil’s Cave (*p* =0.619) or Amis (*p* =0.558) as the proxy of East Asian, suggesting that the East Asian ancestry of Jōmon may phylogenetically be basal to the mainstream northern or southern East Asians. By contrast, the East Asian ancestry of Tibetan, which is probably related to the Tibeto-Burman dissemination in Tibetan Plateau, can be represented by Devil’s Cave (67.7 ± 2.0%, *p* = 0.178) rather than Amis (*p* = 1.24×10^-4^), consistent with the North China origin of the Sino-Tibetan language family^34,35^.

Fitting the scenario of the expansion of Neolithic farmers in southern China, southern East Asian lineage represented by Amis predominates in Hàn (69.3 ± 2.8%), other populations in southern China (69.2–78.6%), and populations in MSEA (54.3–76.7%). Hàn and Tibeto-Burman-speaking populations (Nàxī, Yí, and Tǔjiā) tend to have more Devil’s Cave ancestry (23.5–30.7%) than their neighbors, consistent with our previous inference. Compared with isolated Austroasiatics, Kinh and Thai/Khmer respectively harbor additional Devil’s Cave (18.7 ± 7.5%) and Namazga (8.4 ± 1.0% for Thai, 3.7 ± 1.1% for Khmer) ancestry, likely reflecting immigration from Hàn Chinese and South Asians in the historical period, accordingly. Apart from MSEA, Austroasiatic-speaking Kharia in Central India is also estimated with Ami-like component (18.5 ± 3.0%), indicating a southern East Asian origin of the Muṇḍā branch.

Although the Devil’s Cave ancestry is generally the predominant East Asian lineage in North Asia and adjacent areas, there is an intriguing discrepancy between the eastern [Korean, Japanese, Tungusic (except northernmost Oroqen), and Mongolic (except westernmost Kalmyk) speakers] and the western part [West Xiōngnú (∼2,150 BP)^36^, Tiānshān Hun (∼1,500 BP)^36^, Turkic-speaking Karakhanid (∼1,000 BP)^36^ and Tuva, and Kalmyk]. Whereas the East Asian ancestry of populations in the western part has entirely belonged to the Devil’s Cave lineage till now, populations in the eastern part have received the genomic influence from an Amis-related lineage (17.4–52.1%) posterior to the presence of the Devil’s Cave population roughly in the same region (∼7,600 BP)^12^. Analogically, archaeological record has documented the transmission of wet-rice cultivation from coastal China (Shāndōng and/or Liáoníng Peninsula) to Northeast Asia, notably the Korean Peninsula (Mumun pottery period, since ∼3,500 BP) and the Japanese archipelago (Yayoi period, since ∼2,900 BP)^2^. Especially for Japanese, the Austronesian-related linguistic influence in Japanese^37^ may indicate a potential contact between the Proto-Japonic speakers and population(s) affiliating to the coastal lineage. Thus, our results imply that a southern-East-Asian-related lineage could be arguably associated with the dispersal of wet-rice agriculture in Northeast Asia at least to some extent.

## DISCUSSION

In this study, we discover an explicit genomic pattern regarding Hmong-Mien speakers and other southern East Asians, which has been established in Neolithic and is still detectable in present-day populations. After the separation from the northern East Asians, southern East Asians further divided into two lineages genetically, concordant with archaeological and linguistic evidence. While the inland lineage is associated with the farming dispersal from the middle Yangtze and the dissemination of the Austroasiatic and Hmong-Mien language family, the coastal lineage is more correlated with the agriculture expansion out of Yangtze Delta and the diffusion of the Austronesian and (at least partially) Kra-Dai language family. Especially for Hmong-Mien speakers, the dispersal from inland southern China and subsequent genetic contact with neighboring populations (particularly Tibeto-Burman-and Kra-Dai-speaking ones) majorly form the demographic history of Hmong-Mien-speaking populations.

It has been long debated how the Neolithic cultures relate to major language families in East Asian, and several assumptions have been proposed, notably the connection between the Dàxī culture and Proto-Hmong-Mien^30^, the Dàwènkǒu culture and Pre-Austronesian^30^, as well as the Yǎngsháo^34^/Císhān^35^ culture and Proto-Sino-Tibetan. Even though the ancient genomic data from the Neolithic farming heartlands in China have not been sufficiently reported yet, we still recover a general demographic pattern using both the genetic diversity preserved in present-day populations and ancient samples from peripheral regions of East Asia, which can be served as a foothold for further archaeogenetic studies on the demographic dynamic of the southern East Asia. Although we observe a good matching among the evidence from genetics, linguistics, and archaeology, we still cautiously refer the proposed ancestral populations to an approximate geographical range instead of determined archaeological sites. Thus, we predict further studies on directly sampled ancient genomes from these sites will shed new light on resolving a clearer genetic landscape of this region, especially the demographic history since the Neolithic revolution.

## METHOD

### Samples

Totally, we collected 903 Y-chromosomal samples [including 458 published^38^ samples of 17 loci Y-chromosomal Short Tandem Repeat (Y-STR)] from healthy males, 624 mitochondrial samples, and 415 autosomal samples from unrelated volunteers in Húnán, China. All the samples are either blood or saliva. All the participants had signed the consent forms before joining the study. The study was approved by the ethics committee for biological research at the School of Life Science at Fudan University.

### Molecular Methods

We used the DP-318 Kit (Tiangen Biotechnology, Beijing) to extract the genomic DNA with the manufacturer’s protocol. We used AmpFlSTR® YFiler™ PCR Amplification Kit (Applied Biosystems, Foster City, CA, USA) to amplify and genotype 17 Y-STR loci (DYS19, DYS389a/b, DYS390, DYS391, DYS392, DYS393, DYS437, DYS438, DYS439, DYS448, DYS456, DYS458, DYS635, Y-GATA H4, and DYS385a/b) of all Y-chromosomal samples. We used the ABI 3730 and ABI 3130 Genetic Analyzers (Applied Biosystems) to analyze the amplification products. We used Genscan v. 3.7 and Genotyper v. 3.7 (Applied Biosystems) to analyze electrophoresis results. We selected 936 Y-chromosomal single nucleotide polymorphisms (Y-SNPs) for genotyping, according to Y-haplogroup polymorphism in the East Asian populations. Then we designed single-stranded nucleic acid bait libraries (Roche Nimblegen, Pleasanton, CA, USA) to capture these Y-SNPs. We sent ∼500 of our Y-chromosomal samples for next-generation sequencing on the Illumina HiSeq2000 platform (Illumina, San Diego, CA, USA) and assigned haplogroups according to their Y-SNP results. For other Y-chromosomal samples, the haplogroups were inferred by Y-STR comparison. Other details of our next-generation sequencing process are identical to those previously described^18^. All 624 mitochondrial samples and 415 autosomal samples were extracted and performed similar processes of next-generation sequencing applied for Y-chromosomal samples to attain complete mitochondrial DNA sequences and selected autosomal single nucleotide polymorphisms (SNPs), respectively. Mitochondrial sequences were aligned to the Cambridge Reference Sequence (rCRS) by Kalign^39^ and manually edited by BioEdit 7.0.4.1^40^, with haplogroups assigned in accordance with *phylotree* Build 17^41^.

### Genomic dataset assembly

We used three distinct datasets in population genetic analysis: (1) a dataset for all the 415 newly reported samples from Central China, covering 9,752 SNPs and referred as “10k dataset”; (2) a dataset based on Human Origin (HO) dataset from Reich Lab (https://reich.hms.harvard.edu/datasets) ^42,43^ and combining other ancient and modern samples^3,4,12,17,32,36,44-56^, covering 590,388 SNPs, and referred as “600k dataset”; (3) a dataset based on Simons Genome Diversity Project^17^, combining other ancient and modern samples^3,4,12,32-46^, covering 1,233,013 SNPs, and referred as “1240k dataset”.

### Principle Component Analysis (PCA)

We performed PCA in *smartpca* program of EIGENSOFT^11^ with parameters as default parameters and killr2: YES, r2thresh: 0.1, r2genlim: 0.02, lsqproject: YES, shrinkmode: YES. We used 10k dataset for PCA of population in Húnán, and a combination of 10k and 1240k dataset for East-Asia-wide PCA (leaving 4,911 SNPs). For East-Asia-wide PCA, principle components are constructed by modern samples with high-coverage ancient samples [Shamanka_EN (DA249)^32^, a Devil’s Cave individual^12^ (NEO240), Chokhopani^45^, and Laos_LN_BA^3^ (LA364)] being projected.

### Model-based clustering

We first applied PLINK^57^ with the parameter indep-pairwise 50 5 0.2 to prune out the Linkage Disequilibrium (LD), yielding 3995 SNPs (10k dataset). Then, we performed the ADMIXTURE^14^ from K = 2 to K = 10. We find the likelihood increasing and when K≥4, there is a heterogeneous composition within Pahng, which is inconsistent with the result from PCA and *f*_*3*_ statistics. Therefore, we assume the best modeling reaches when K = 3.

### *f*-statistics

We utilized the *qp3Pop* and *qpDstat* program of ADMIXTOOLS^16^ to calculate *f*_*3*_-and *D*-statistics with default parameters. Based on the definition in Raghavan *et al.*, 2014^52^, we calculated the outgroup *f*_*3*_(Mbuti; Hmong, X) to measure the shared drift between Hmong and Eurasian populations, using the 1240k dataset. Based on the result of outgroup *f*_*3*_-statistics, we computed the main *D* (X_1_, Mbuti; Hmong, X_2_), where X_1_ = Eurasia-wide populations and X_2_ = populations shared the most genetic drift with Hmong with additional *D*-statistics.

### TreeMix analysis

We used the TreeMix^15^ to generate the admixture graph, with k = 10 and *noss* for the 10k dataset, and k = 1000 and *noss* for the 600k dataset. We set migration events from m = 1 to m = 10 and reported the result of m = 4.

### Coalescence time estimation

In total, 93 complete mitochondrial sequences were selected for phylogenetic analysis of haplogroup B5, including 52 sequences from this study and 41 sequences [33 of B5, 1 of B4’5*^58^ (Tiányuán), 1 of B4, and 6 of outgroup of B4’5] from phylotree Build 17^41^. The mutations of C indels 309.1C(C)-315.1C, AC indels at 515-522, C indels at 956-960, 3107, 8281-8289, A16182c, A16183c, 16189 and 16193.1C(C) were excluded from phylogeny reconstruction, manually performed using BioEdit 7.0.4.1^40^. We applied ARB^59^ to generate the phylogeny using the Maximum Parsimony (MP) method. Bayesian time estimation for the most recent common ancestor (TMRCA) of B5 and its subclades were performed by BEAST v1.10.0^60^. The monophyletic option was set according to the previously generated MP phylogeny. The age of 40.0 kya for Tiányuán belonging to mitochondrial haplogroup B4’5* was chosen for time calibration^58^. For substitution model, HKY model was chosen with estimated base frequencies and Site Heterogeneity Model was set to eight Gamma categories. Strict clock model was set, and Bayesian skyline model was applied as prior^61^. The run was conducted 10 million iterations with sampling every 1000 steps, with the first 10% burnt in. All the effective sample sizes (ESS) were higher than 200. We applied similar procedure to the full-sequencing Y-chromosomal data for Bayesian TMRCA estimation.

### Admixture graph fitting by *qpGraph*

We used the *qpGraph* program of ADMIXTOOLS^16^ to create the admixture graph based on the result of *D*-statistics, with the default parameters and useallsnps: YES, outpop: NULL. We used 600k dataset to fit the admixture graph, leaving 496,352 SNPs. We used the backbone graph described in McColl *et al.*, 2018^3^ as the initial graph. To avoid the case of a “basal” admixture, only Mbuti and Eurasian samples without significant Denisovan ancestry retained in the graph. Since all the Eurasian samples in the graph (except Ust’-Ishim and Tiányuán) do not harbor significantly excess Neanderthal ancestry compared with each other^3^, we suppose that the interference of Neanderthal ancestry can only affect fitness between Mbuti and Eurasian samples, and Tiányuán or Ust’-Ishim with the others. Thus, such interference cannot significantly bias the main result of our *qpGraph* fitting.

### Inference of mixture proportions

We used the *qpWave*^*62*^ and *qpAdm*^31^ program of ADMIXTOOLS to estimate the fitness and proportion of admixture components. We used the Devil’s Cave samples^12^, Amis, the Laotian Hòabìnhian individual (LA368)^3^, Afanasievo^56^, Namazga^32^, and Kolyma^12^ as reference populations. We used Mbuti, Peștera cu Oase1^53^, Ust’-Ishim^48^, Tiányuán^47^, Kostenki^49^, Yana_UP^12^, Naṭūfian_N^43^, Anatolia_N^54^, Motala_HG^54,55^, Önge^17^, Papuan, Eskimo, and Mixe as outgroups.

### Inbreeding coefficient F_ST_

Using 10k dataset, we performed *smartpca* program of EIGENSOFT^11^ to estimate pairwise F_ST_ with default parameters and fstonly: YES.

## Abbreviations

We use the following abbreviated labels addressing populations throughout the paper:

AAA: Ancestral Austroasiatic;
AAN: Ancestral Austronesian;
AHM: Ancestral Hmong-Mien;
AKD: Ancestral Kra-Dai;
South_EA: southern East Asian;
North_EA: northern East Asians;
First_MSEA: First Mainland Southeast Asian;
Second_MSEA: Second Mainland Southeast Asian;
Insular_AN: Insular Austronesian;
SEAH: southern East Asian ancestry of Hàn;
Aus: Australasian;
UP: Upper Pleistocene;
N: Neolithic;
EN: Early Neolithic;
LN: Late Neolithic;
BA: Bronze Age;
IA: Iron Age;
HG: hunter-gatherer.

---

**Extended Data Figure 1.**
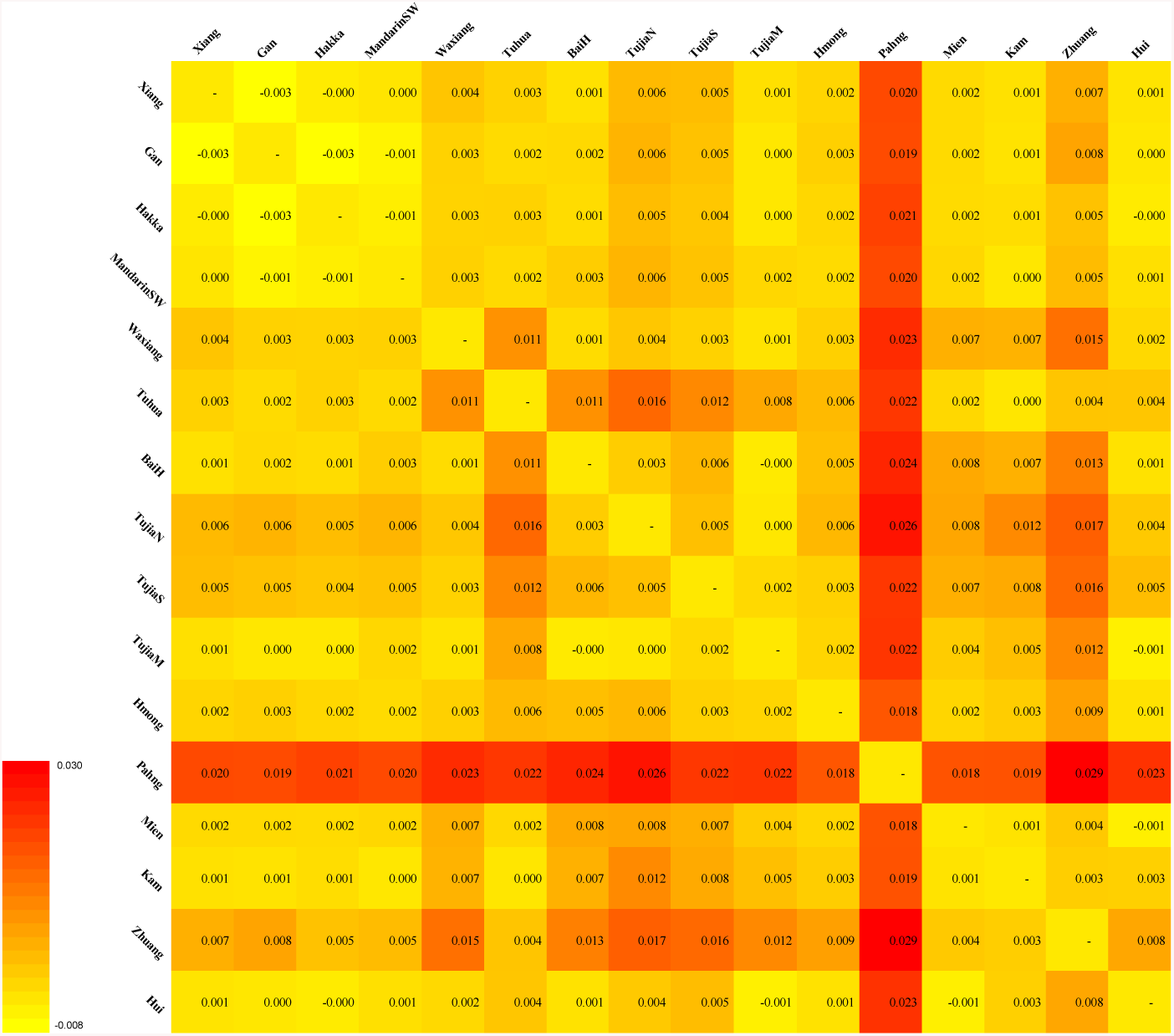
Pairwise F_ST_ of populations in Húnán.

**Extended Data Figure 2.**
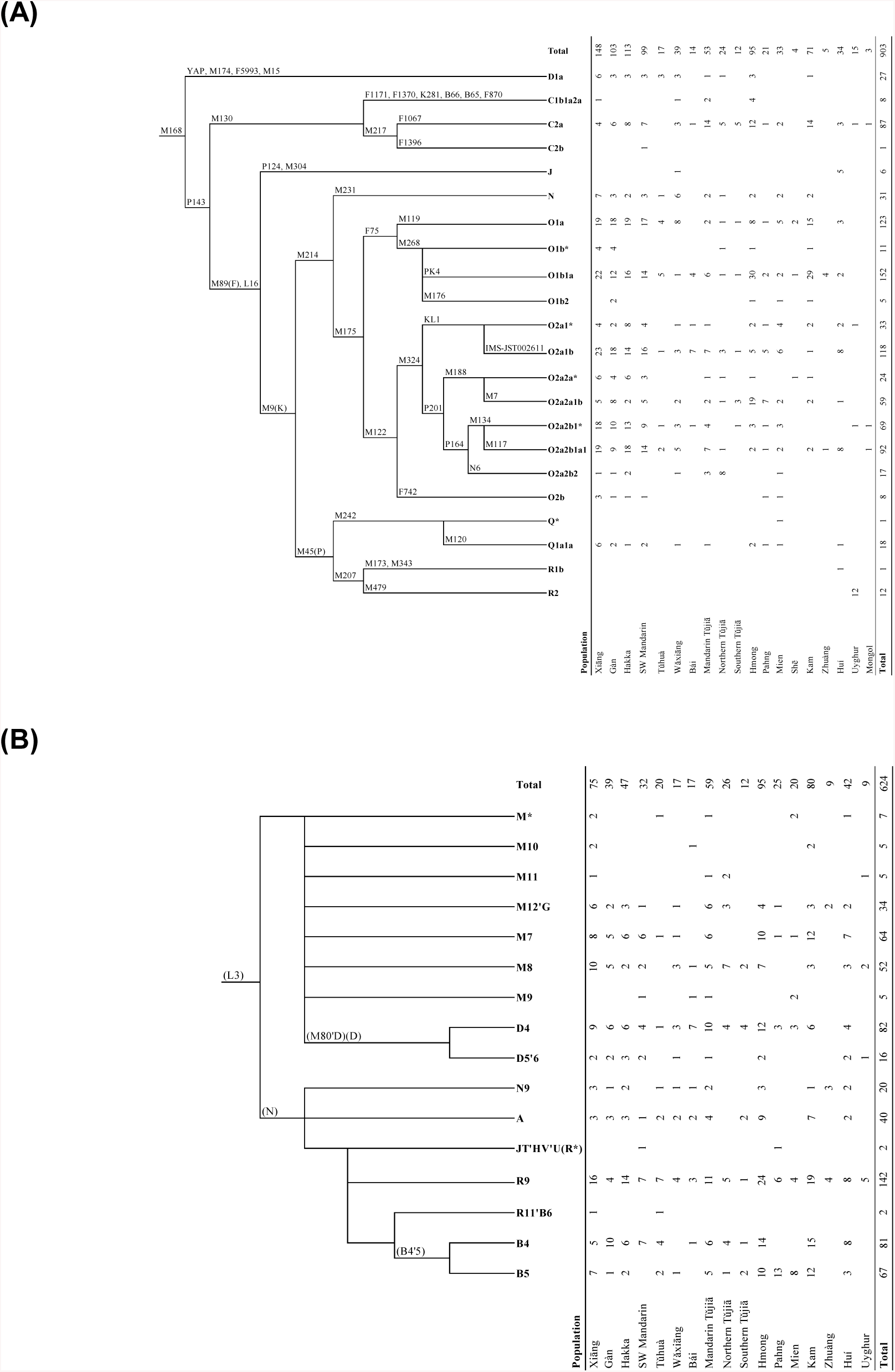
Phylogenies and distribution of (A) Y-chromosomal and (B) mitochondrial haplogroups in populations in Húnán.

**Extended Data Figure 3.**
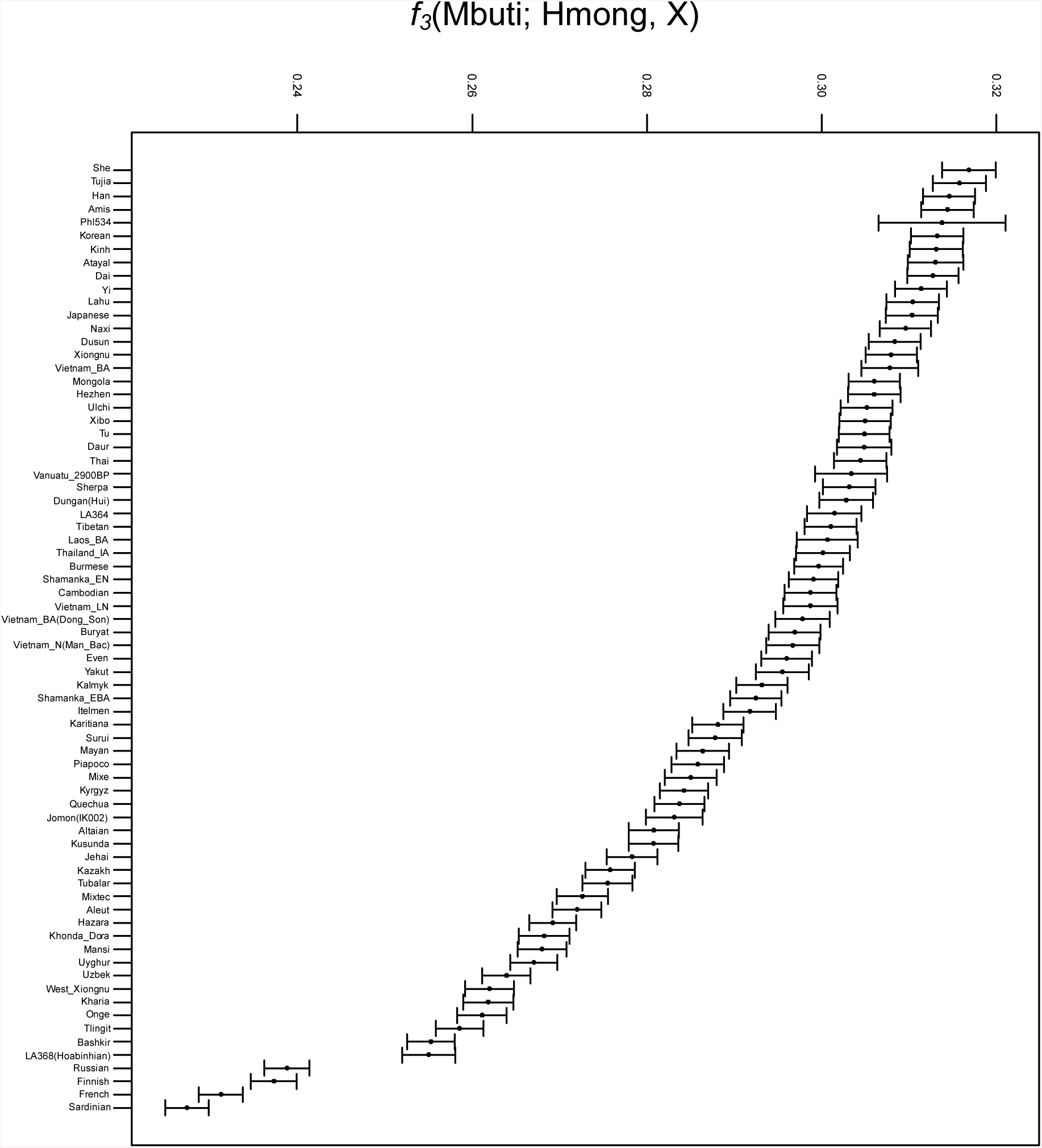
Outgroup *f*_*3*_-statistics measuring shared drift between Hmong and other Eurasian populations. Mbuti is used as the outgroup.

**Extended Data Figure 4.**
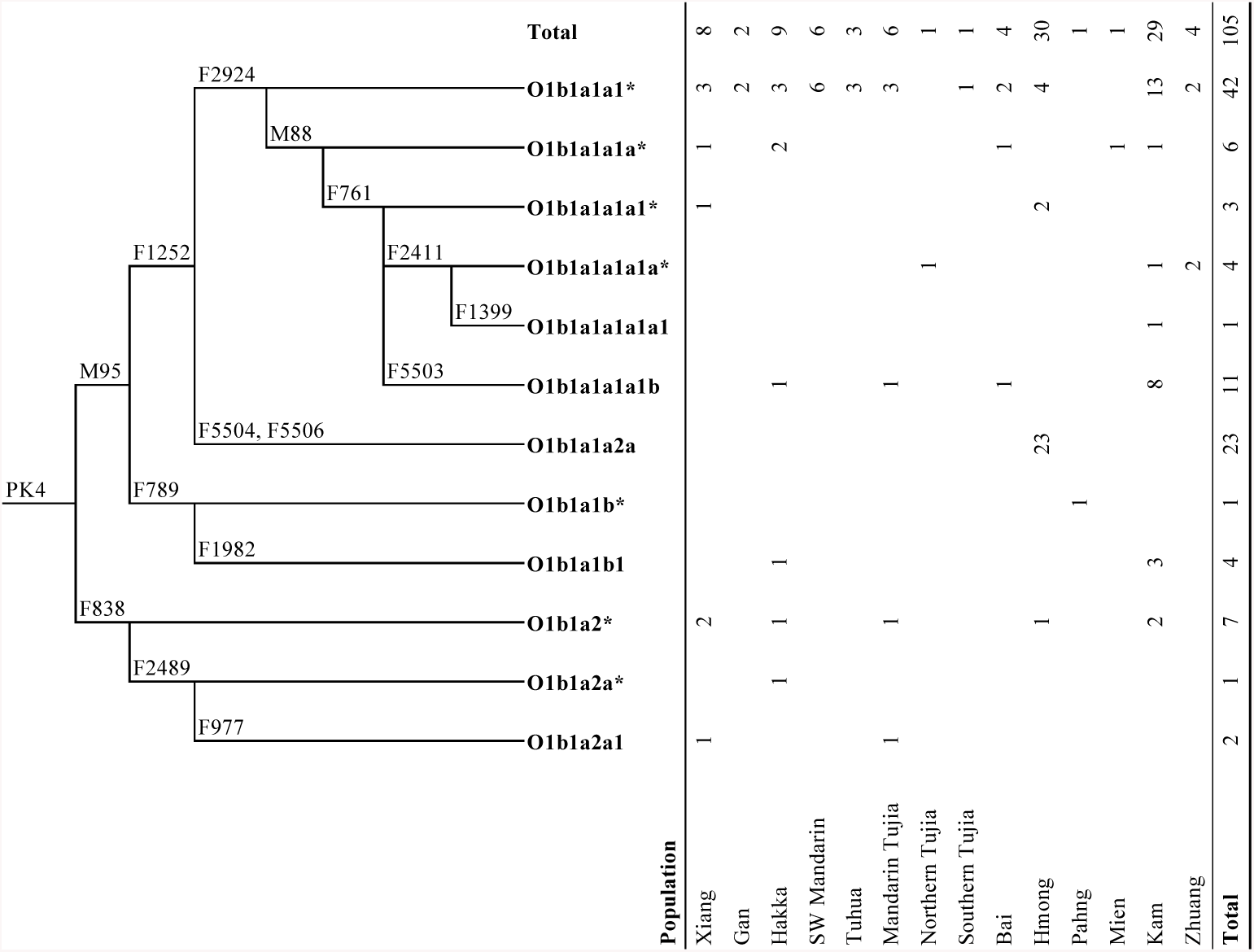
Distribution of subgroups of Y-haplogroup O1b1a-PK4 in Húnán.

**Extended Data Table 1.** Autosomal sample size of populations in Hunan.

| Category | Population | Sample size |
| --- | --- | --- |
| Sinitic | Xiāng | 25 |
|  | Gàn | 28 |
|  | Hakka | 47 |
|  | MandarinSW | 6 |
|  | Huí | 4 |
|  | Tǔhuà | 12 |
|  | Wǎxiāng | 15 |
|  | Báih | 14 |
| Tibeto-Burman | TǔjiāN | 23 |
|  | TǔjiāS | 12 |
|  | TǔjiāM | 46 |
| Hmong-Mien | Hmong | 79 |
|  | Pahng | 16 |
|  | Mien | 11 |
| Kra-Dai | Kam | 72 |
|  | Zhuàng | 5 |
| Total |  | 415 |

**Extended Data Table 2.** *D*-statistics with only transversions for tetrads including ancient samples in Table 2.

| Populations | *D* (X, Mbuti;<br>Hmong, Tujia) |  | *D* (X, Mbuti;<br>Hmong, Han) |  | *D* (X, Mbuti;<br>Hmong, She) |  | *D* (X, Mbuti;<br>Hmong, Dai) |  | *D* (X, Mbuti;<br>Hmong, Amis) |  |
| --- | --- | --- | --- | --- | --- | --- | --- | --- | --- | --- |
|  | mean | Z-score | mean | Z-score | mean | Z-score | mean | Z-score | mean | Z-score |
| *DevilsCave_N* | -0.0027 | -1.347 | -0.0050 | -3.028 | -0.0026 | -1.157 | 0.0105 | 4.998 | 0.0050 | 1.910 |
| *Shamanka_EN* | -0.0022 | -1.332 | -0.0027 | -2.006 | -0.0015 | -0.833 | 0.0106 | 6.201 | 0.0071 | 3.355 |
| *Lokomotiv_EN* | -0.0031 | -1.467 | -0.0023 | -1.302 | -0.0028 | -1.175 | 0.0109 | 4.712 | 0.0061 | 2.217 |
| *Ust'-Ida_LN* | -0.0043 | -2.008 | -0.0030 | -1.720 | -0.0025 | -1.087 | 0.0087 | 4.059 | 0.0011 | 0.407 |
| *Shamanka_EBA* | -0.0037 | -1.819 | -0.0041 | -2.479 | -0.0029 | -1.334 | 0.0094 | 4.338 | 0.0031 | 1.212 |
| *Okunevo_BA* | -0.0037 | -2.170 | -0.0033 | -2.403 | -0.0046 | -2.446 | 0.0033 | 1.863 | 0.0002 | 0.115 |
| *Xiōngnú* | -0.0028 | -1.356 | -0.0044 | -2.501 | -0.0047 | -2.083 | 0.0071 | 3.202 | 0.0005 | 0.170 |
| *Xiōngnú_West* | -0.0005 | -0.210 | -0.0004 | -0.212 | -0.0019 | -0.784 | 0.0057 | 2.453 | -0.0005 | -0.196 |
| *Chokhopani* | -0.0015 | -0.618 | -0.0012 | -0.598 | -0.0034 | -1.335 | 0.0129 | 5.149 | 0.0094 | 3.128 |
| *Laos_LN_BA (LA364)* | 0.0016 | 0.559 | 0.0030 | 1.334 | -0.0009 | -0.312 | -0.0103 | -3.753 | -0.0047 | -1.456 |
| *Laos_BA* | 0.0100 | 2.430 | 0.0062 | 1.925 | -0.0027 | -0.614 | -0.0062 | -1.524 | -0.0087 | -1.750 |
| *Vietnam_N (Mán Bạc)* | 0.0056 | 1.580 | 0.0067 | 2.213 | 0.0031 | 0.774 | -0.0062 | -1.672 | 0.0004 | 0.087 |
| *Thailand_LN_IA (Ban Chiang)* | 0.0081 | 1.146 | 0.0095 | 1.671 | 0.0002 | 0.033 | 0.0003 | 0.041 | 0.0021 | 0.245 |
| *Vietnam_LN* | -0.0035 | -1.008 | 0.0016 | 0.573 | -0.0063 | -1.712 | -0.0055 | -1.523 | -0.0096 | -2.245 |
| *Vietnam_BA (Đồng Sơn)* | 0.0041 | 1.227 | 0.0028 | 0.977 | -0.0051 | -1.347 | -0.0049 | -1.351 | -0.0064 | -1.470 |
| *Vietnam_BA (Núi Nạp)* | 0.0005 | 0.144 | -0.0009 | -0.288 | -0.0062 | -1.517 | -0.0061 | -1.596 | -0.0102 | -2.297 |
| *Anzick* | 0.0005 | 0.224 | -0.0025 | -1.221 | -0.0030 | -1.125 | 0.0078 | 3.040 | 0.0024 | 0.785 |
| *Hòabìnhian_Laos (LA368)* | -0.0012 | -0.351 | -0.0024 | -0.886 | -0.0091 | -2.485 | -0.0028 | -0.774 | -0.0113 | -2.698 |
| *Tiányuán* | -0.0042 | -1.606 | -0.0037 | -1.714 | -0.0047 | -1.730 | -0.0071 | -2.556 | -0.0060 | -1.849 |
| *Jōmon* | -0.0034 | -1.326 | -0.0003 | -0.144 | -0.0036 | -1.300 | 0.0057 | 2.073 | -0.0118 | -3.525 |

**Extended Data Table 3.**
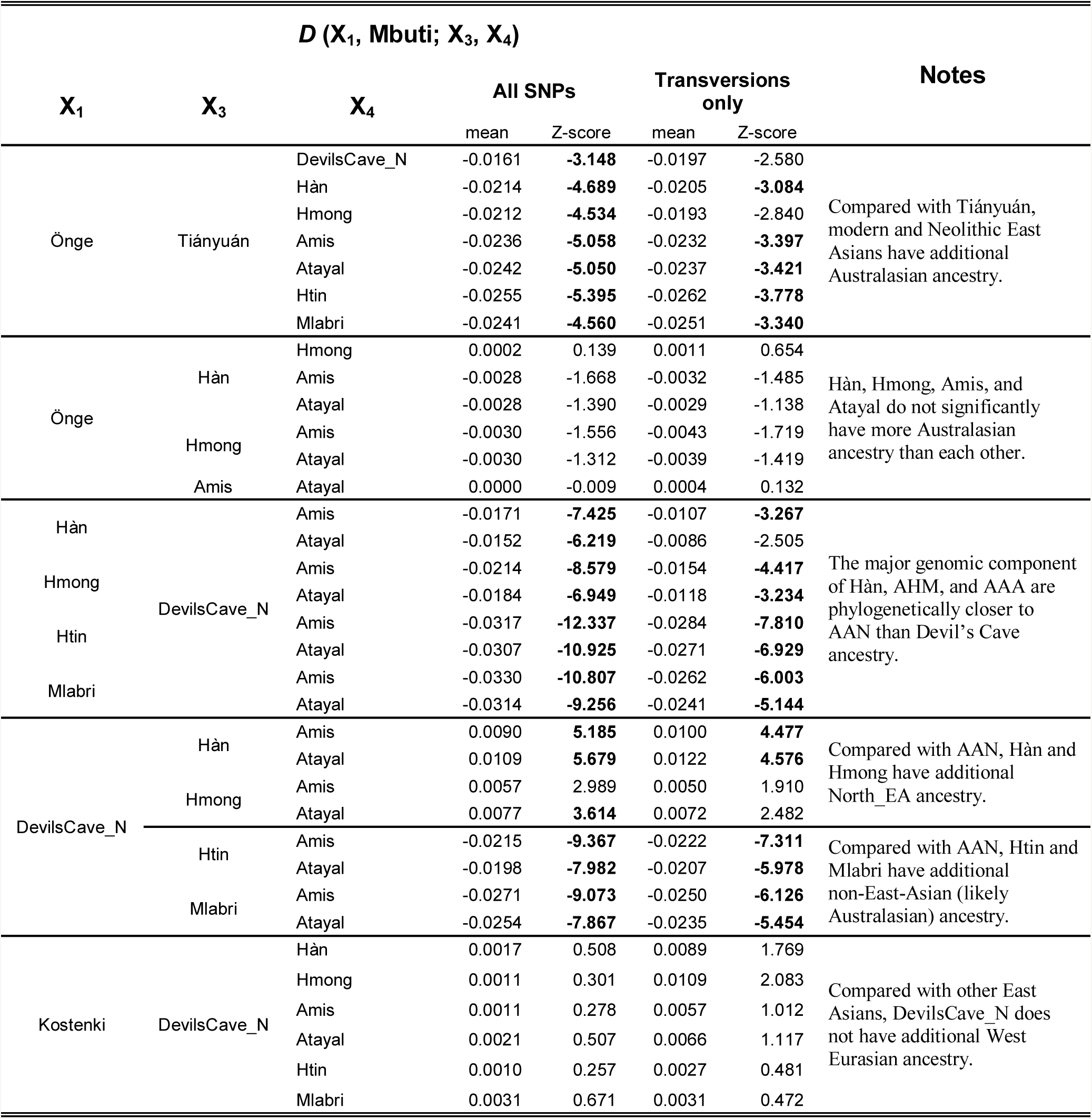
Additional *D*-statistics.

**Extended Data Table 4.** Original *qpAdm* results.

| Test Population | Feasible | Best Model | p-value | Amis |  | Devil's_Cave |  | Hòabinhian |  | Afanasievo |  | Namazga |  | Kolyma |  |
| --- | --- | --- | --- | --- | --- | --- | --- | --- | --- | --- | --- | --- | --- | --- | --- |
|  |  |  |  | mean | str_err | mean | str_err | mean | str_err | mean | str_err | mean | str_err | mean | str_err |
| Hàn | Yes | Yes | 8.912E-01 | 0.6927 | 0.028 | 0.3073 | 0.028 | NA | NA | NA | NA | NA | NA | NA | NA |
| Huí | Yes | Yes | 8.526E-01 | 0.4480 | 0.067 | 0.3925 | 0.060 | NA | NA | 0.1062 | 0.044 | 0.0533 | 0.046 | NA | NA |
| Xiōngnú | Yes | Yes | 5.427E-01 | 0.5014 | 0.070 | 0.4567 | 0.064 | NA | NA | 0.0418 | 0.013 | NA | NA | NA | NA |
| Tǔjiǎ | Yes | Yes | 9.913E-01 | 0.7063 | 0.034 | 0.2937 | 0.034 | NA | NA | NA | NA | NA | NA | NA | NA |
| Nàxī | Yes | Yes | 9.118E-01 | 0.7649 | 0.035 | 0.2351 | 0.035 | NA | NA | NA | NA | NA | NA | NA | NA |
| Yí | Yes | Yes | 9.425E-01 | 0.7326 | 0.034 | 0.2674 | 0.034 | NA | NA | NA | NA | NA | NA | NA | NA |
| Tibetan | Yes | Yes | 1.782E-01 | NA | NA | 0.6775 | 0.020 | 0.3225 | 0.020 | NA | NA | NA | NA | NA | NA |
|  | No | No | 1.235E-04 | 1.0005 | 0.033 | NA | NA | -0.0005 | 0.033 | NA | NA | NA | NA | NA | NA |
|  | No | No | 1.635E-01 | -0.1374 | 0.420 | 0.7712 | 0.286 | 0.3662 | 0.136 | NA | NA | NA | NA | NA | NA |
| Lāhù | Yes | Yes | 7.074E-01 | 0.6916 | 0.150 | 0.1930 | 0.102 | 0.1154 | 0.050 | NA | NA | NA | NA | NA | NA |
| Korean | Yes | Yes | 6.460E-01 | 0.4488 | 0.041 | 0.5512 | 0.041 | NA | NA | NA | NA | NA | NA | NA | NA |
| Japanese | Yes | Yes | 7.293E-01 | 0.5208 | 0.032 | 0.4792 | 0.032 | NA | NA | NA | NA | NA | NA | NA | NA |
| Jōmon | Yes | Yes | 6.185E-01 | NA | NA | 0.4191 | 0.027 | 0.5809 | 0.027 | NA | NA | NA | NA | NA | NA |
|  | Yes | No | 5.582E-01 | 0.6076 | 0.039 | NA | NA | 0.3924 | 0.039 | NA | NA | NA | NA | NA | NA |
|  | No | No | 6.108E-01 | 0.3535 | 0.334 | 0.1798 | 0.230 | 0.4667 | 0.109 | NA | NA | NA | NA | NA | NA |
| Xībó | Yes | Yes | 9.090E-01 | 0.3519 | 0.046 | 0.5988 | 0.043 | NA | NA | 0.0493 | 0.009 | NA | NA | NA | NA |
| Hezhen | Yes | Yes | 4.004E-01 | 0.1845 | 0.054 | 0.7954 | 0.049 | NA | NA | 0.0201 | 0.011 | NA | NA | NA | NA |
| Oroqen | No | No | 4.657E-01 | -0.0044 | 0.063 | 0.9651 | 0.057 | NA | NA | 0.0393 | 0.013 | NA | NA | NA | NA |
|  | Yes | No | 5.833E-02 | 0.0671 | 0.054 | 0.9329 | 0.054 | NA | NA | NA | NA | NA | NA | NA | NA |
|  | Yes | Yes | 5.631E-01 | NA | NA | 0.9615 | 0.011 | NA | NA | 0.0385 | 0.011 | NA | NA | NA | NA |
| Monguor | Yes | Yes | 8.242E-01 | 0.5151 | 0.043 | 0.3927 | 0.038 | NA | NA | NA | NA | 0.0922 | 0.009 | NA | NA |
| Mongol | Yes | Yes | 9.025E-02 | 0.4047 | 0.049 | 0.5309 | 0.045 | NA | NA | 0.0645 | 0.010 | NA | NA | NA | NA |
| Daur | Yes | Yes | 3.295E-01 | 0.1741 | 0.053 | 0.7751 | 0.049 | NA | NA | 0.0508 | 0.011 | NA | NA | NA | NA |
| Shamanka_EN | Yes | Yes | 6.520E-01 | NA | NA | 0.6796 | 0.051 | NA | NA | 0.0176 | 0.011 | NA | NA | 0.3028 | 0.053 |
|  | No | No | 5.227E-01 | 0.0190 | 0.094 | 0.6538 | 0.152 | NA |  | 0.0154 | 0.017 | NA | NA | 0.3117 | 0.081 |
| Xiongnu_West | No | No | 1.252E-01 | 0.0124 | 0.083 | 0.3733 | 0.076 | NA | NA | 0.6143 | 0.017 | NA | NA | NA | NA |
|  | Yes | Yes | 1.734E-01 | NA | NA | 0.3839 | 0.015 | NA | NA | 0.6161 | 0.015 | NA | NA | NA | NA |
| Hun_Tianshan | Yes | Yes | 3.000E-01 | NA | NA | 0.2392 | 0.008 | NA | NA | 0.6536 | 0.044 | 0.1072 | 0.044 | NA | NA |
|  | No | No | 3.320E-01 | -0.0805 | 0.055 | 0.3107 | 0.050 | NA | NA | 0.6272 | 0.046 | 0.1426 | 0.048 | NA | NA |
| Karakhanid | Yes | Yes | 2.045E-01 | NA | NA | 0.4271 | 0.012 | NA | NA | 0.3848 | 0.052 | 0.1881 | 0.051 | NA | NA |
|  | No | No | 1.336E-01 | 0.0071 | 0.087 | 0.4211 | 0.080 | NA | NA | 0.3888 | 0.060 | 0.1829 | 0.064 | NA | NA |
| Tuva | Yes | Yes | 3.123E-01 | NA | NA | 0.7920 | 0.010 | NA | NA | 0.2080 | 0.010 | NA | NA | NA | NA |
|  | No | No | 3.623E-01 | -0.0694 | 0.058 | 0.8538 | 0.053 | NA | NA | 0.2156 | 0.012 | NA | NA | NA | NA |
| Kalmyk | Yes | Yes | 4.417E-01 | NA | NA | 0.7510 | 0.010 | NA | NA | 0.2490 | 0.010 | NA | NA | NA | NA |
|  | No | No | 3.545E-01 | 0.0051 | 0.051 | 0.7464 | 0.047 | NA | NA | 0.2485 | 0.011 | NA | NA | NA | NA |
| Hmong | Yes | Yes | 9.715E-01 | 0.7861 | 0.032 | 0.2139 | 0.032 | NA | NA | NA | NA | NA | NA | NA | NA |
| She | Yes | Yes | 7.857E-01 | 0.7397 | 0.035 | 0.2603 | 0.035 | NA | NA | NA | NA | NA | NA | NA | NA |
| Kinh_all | Yes | Yes | 9.547E-01 | 0.7117 | 0.109 | 0.1874 | 0.075 | 0.1010 | 0.036 | NA | NA | NA | NA | NA | NA |
| Dai | Yes | Yes | 6.580E-01 | 0.7296 | 0.164 | 0.1759 | 0.112 | 0.0945 | 0.054 | NA | NA | NA | NA | NA | NA |
| Thai | No | No | 5.396E-01 | 0.6492 | 0.169 | 0.0696 | 0.115 | 0.1999 | 0.063 | NA | NA | 0.0813 | 0.012 | NA | NA |
|  | Yes | Yes | 6.916E-01 | 0.7504 | 0.017 | NA | NA | 0.1654 | 0.021 | NA | NA | 0.0843 | 0.010 | NA | NA |
| Khmer | Yes | Yes | 9.877E-01 | 0.5425 | 0.127 | 0.1192 | 0.087 | 0.3009 | 0.047 | NA | NA | 0.0374 | 0.011 | NA | NA |
| Htin | No | No | 8.426E-01 | 0.6729 | 0.150 | 0.0657 | 0.105 | 0.2614 | 0.048 | NA | NA | NA | NA | NA | NA |
|  | Yes | Yes | 8.213E-01 | 0.7669 | 0.021 | NA | NA | 0.2331 | 0.021 | NA | NA | NA | NA | NA | NA |
| Mlabri | Yes | Yes | 4.001E-01 | 0.7252 | 0.029 | NA | NA | 0.2748 | 0.029 | NA | NA | NA | NA | NA | NA |
|  | No | No | 3.397E-01 | 0.5970 | 0.293 | 0.0901 | 0.204 | 0.3130 | 0.093 | NA | NA | NA | NA | NA | NA |
| Kharia | Yes | Yes | 4.649E-01 | 0.1849 | 0.030 | NA | 0.041 | 0.6160 | 0.022 | NA | NA | 0.1991 | NA | NA | NA |
|  | No | No | 3.924E-01 | 0.4584 | 0.338 | -0.2007 | 0.244 | 0.5305 | 0.116 | NA | NA | 0.2118 | 0.027 | NA | NA |

