## Supplementary Note 1 for "Inland-coastal bifurcation of southern East Asians revealed by Hmong-Mien genomic history"

### Formally inferring and examining phylogenetic models from actual *D*-statistics

#### 1.1. Motivation and convention

A single $F_4$ (Test$_1$, Outgroup; Test$_2$, Test$_3$)[1] can determine an overall phylogeny for the three test populations. However, if we are interested in the phylogenetic relationship among a series of ghost populations (*e.g.*, the 'Basal Eurasian'[2]), since the ghost populations do not exist in unmixed forms, we cannot directly infer the phylogeny merely from the direction of the $f_4$-statistics. Admixture graph fitting by *qpGraph*[1] is a helpful way to settle it. However, it can be difficult to find the best fitting when the test populations have a complex demographic history that is hard to be globally modeled.

Inspired by the definition of 'shared drift' for *f*-statistics[1], we focus on a particular case where Test$_2$ and Test$_3$ do not have significant excess deep ancestry than each other. Given that, the deep ancestry of Test$_1$ cannot affect the result of $F_4$ (Test$_1$, Outgroup; Test$_2$, Test$_3$), because $F_4$ (deep ancestry, Outgroup; Test$_2$, Test$_3$) is supposed to be zero. Therefore, we can determine the phylogeny among the shared and non-deep ancestry of Test$_1$ and different components of Test$_2$ and Test$_3$. In the following part, we infer the phylogeny of the ghost populations from the *D*-statistics involving First Mainland Southeast Asians and Insular Austronesians and examine the hypothesized phylogeny against the actual *D*-statistics of Second Mainland Southeast Asians. We deem the theoretical $F_4$ significantly deviate from zero if $|Z| > 2.58$ for corresponding *D*-statistics (two-tailed *p*-value < 0.01).

We use the following abbreviations in the following part: AAA, Ancestral Austroasiatic; AAN, Ancestral Austronesian; AHM, Ancestral Hmong-Mien; AKD, Ancestral Kra-Dai; Aus, Australasian; DevilsCave, Devil's Cave samples; SEAH, Southern East Asian ancestry of Hàn; First_MSEA, First Mainland Southeast Asian; Insular_AN, Insular Austronesian.

#### 1.2. AAA closer to AHM than AAN (SI Fig. 1.1A)

Since First_MSEA (represented by Htin and Mlabri) has excessive deep Australasian ancestry than AAN (represented by Amis and Atayal, Extended Data Table 3), and AAA and AAN are supposed to symmetrically relate to Australasians, we model Australasian contribution to First_MSEA as

$$\text{First\_MSEA} = \alpha \cdot \text{AAA} + (1-\alpha) \cdot \text{Aus} \ (0 < \alpha < 1) \quad \text{(S 1.1)}$$

Given the idea of 'shared drift', we have

$$F_4(\text{First\_MSEA}, \text{O};\ \text{Hmong}, \text{Amis}) =$$
$$\alpha \cdot F_4(\text{AAA}, \text{O};\ \text{Hmong}, \text{Amis}) + (1-\alpha) \cdot F_4(\text{Aus}, \text{O};\ \text{Hmong}, \text{Amis}) \quad \text{(S 1.2)}$$

Since Hmong and Amis do not significantly have excessive Australasian ancestry than each other (Extended Data Table 3), we have

$$F_4(\text{Aus}, \text{O};\ \text{Hmong}, \text{Amis}) = 0 \quad \text{(S 1.3)}$$

By S 1.2 and 1.3, we have

$$F_4(\text{First\_MSEA}, \text{O};\ \text{Hmong}, \text{Amis}) = \ \alpha \cdot F_4(\text{AAA}, \text{O};\ \text{Hmong}, \text{Amis}) \quad \text{(S 1.4)}$$

By the result in Table 2, we have

$$F_4(\text{First\_MSEA}, \text{O};\ \text{Hmong}, \text{Amis}) = 0 \quad \text{(S 1.5)}$$

By S 1.4 and 1.5, we have

$$F_4(\text{AAA}, \text{O};\ \text{Hmong}, \text{Amis}) = 0 \quad \text{(S 1.6)}$$

Given S 1.6 and the assumption that Amis and Atayal are directly descendent from AAN, we have

$$F_4(\text{AAA}, \text{O};\ \text{Hmong}, \text{AAN}) = 0 \quad \text{(S 1.7)}$$

Since Hmong has an excessive affinity to northern East Asians (especially represented by Devil's Cave samples) than Amis (Table 2), and AHM and AAN are supposed to symmetrically relate to northern East Asians, we model northern East Asian contribution to Hmong as

$$\text{Hmong} = \beta \cdot \text{AHM} + (1-\beta) \cdot \text{DevilsCave}\ (0 < \beta < 1) \quad \text{(S 1.8)}$$

By S 1.8, we have

$$F_4(\text{AAA}, \text{O};\ \text{Hmong}, \text{AAN})$$
$$= \beta \cdot F_4(\text{AAA}, \text{O};\ \text{AHM}, \text{AAN}) + (1-\beta) \cdot F_4(\text{AAA}, \text{O};\ \text{DevilsCave}, \text{AAN}) \quad \text{(S 1.9)}$$

Given that northern East Asian lineage is outgroup to both AAA and AAN, we have

$$F_4\ (\text{AAA}, \text{O};\ \text{DevilsCave}, \text{AAN}) \ll 0 \quad \text{(S 1.10)}$$

By S 1.9 and 1.10, we have

$$F_4(\text{AAA}, \text{O};\ \text{AHM}, \text{AAN}) = -\frac{1-\beta}{\beta} \cdot F_4\ (\text{AAA}, \text{O};\ \text{DevilsCave}, \text{AAN}) \gg 0 \quad \text{(S. 1.11)}$$

Thus, AAA is phylogenetically closer to AHM than AAN. This completes the proof. ■

**1.3. AAN closer to Southern East Asian ancestry of Hàn than AHM** (SI Fig. 1.1B)

Since Hàn has a slightly excessive affinity to northern East Asians than Hmong, we model the ancestral composition of Hàn as

$$\mathrm{Han} = \gamma \cdot \mathrm{SEAH} + (1-\gamma) \cdot \mathrm{DevilsCave}\ (0 < \gamma < 1) \quad \text{(S 1.12)}$$

By S 1.12, we have

$$F_4(\mathrm{Insular\_AN}, \mathrm{O}; \mathrm{Hmong}, \mathrm{Han})$$

$$= \gamma \cdot F_4(\mathrm{Insular\_AN}, \mathrm{O}; \mathrm{Hmong}, \mathrm{SEAH}) + (1-\gamma) \cdot F_4(\mathrm{Insular\_AN}, \mathrm{O};\ \mathrm{Hmong}, \mathrm{DevilsCave}) \quad \text{(S 1.13)}$$

Similarly to the outgroup case of S 1.3, we can neglect the deep ancestry in Insular_AN. Meanwhile, to simplify the modeling, we use AHM to replace Hmong. Given S 1.13, we have

$$F_4(\mathrm{Insular\_AN}, \mathrm{O}; \mathrm{Hmong}, \mathrm{Han})$$

$$= \gamma \cdot F_4(\mathrm{AAN}, \mathrm{O}; \mathrm{AHM}, \mathrm{SEAH}) + (1-\gamma) \cdot F_4(\mathrm{AAN}, \mathrm{O};\ \mathrm{AHM}, \mathrm{DevilsCave}) \quad \text{(S 1.14)}$$

By the result in Table 2, we have

$$F_4(\mathrm{Insular\_AN}, \mathrm{O}; \mathrm{Hmong}, \mathrm{Han}) = 0 \quad \text{(S 1.15)}$$

Since Devil's Cave is outgroup to AAN and AHM, we have

$$F_4(\mathrm{AAN}, \mathrm{O};\ \mathrm{AHM}, \mathrm{DevilsCave}) \gg 0 \quad \text{(S 1.16)}$$

Given S 1.14, 1.15, and 1.16, we have

$$F_4(\mathrm{AAN}, \mathrm{O}; \mathrm{AHM}, \mathrm{SEAH}) = -\frac{1-\gamma}{\gamma} \cdot F_4(\mathrm{AAN}, \mathrm{O};\ \mathrm{AHM}, \mathrm{DevilsCave}) \ll 0 \quad \text{(S 1.17)}$$

Thus, AAN is phylogenetically closer to SEAH than AHM. This completes the proof. ■

**1.4. Checking the dual ancestries of Second_MSEA**

We name the genomic component missing in First_MSEA but existing in Second_MSEA as "Ancestral Kra-Dai" (AKD), and suppose that it is a sister lineage of AAN. Then we have

$$\mathrm{Second\_MSEA} = \delta \cdot \mathrm{First\_MSEA} + (1-\delta) \cdot \mathrm{AKD}\ (0 < \delta < 1) \quad \text{(S 1.18)}$$

By S 1.18, we have

$$F_4(\mathrm{Second\_MSEA}, \mathrm{O}; \mathrm{Hmong}, \mathrm{Han})$$

$$= \delta \cdot F_4(\mathrm{First\_MSEA}, \mathrm{O}; \mathrm{Hmong}, \mathrm{Han}) + (1-\delta) \cdot F_4(\mathrm{AKD}, \mathrm{O}; \mathrm{Hmong}, \mathrm{Han}) \quad \text{(S 1.19)}$$

and

$$F_4(\mathrm{Second\_MSEA}, \mathrm{O}; \mathrm{Hmong}, \mathrm{Amis})$$

$$= \delta \cdot F_4(\text{First\_MSEA}, \text{O}; \text{Hmong}, \text{Amis}) + (1 - \delta) \cdot F_4(\text{AKD}, \text{O}; \text{Hmong}, \text{Amis}) \quad (\text{S } 1.20)$$

Then we examine if our assumption is consistent with the actual *D*-statistics.

##### 1.4.1. Second_MSEA closer to Hmong than Han

Given the result in Table 2, we have

$$F_4(\text{First\_MSEA}, \text{O}; \text{Hmong}, \text{Han}) \gg 0 \quad (\text{S } 1.21)$$

By S 1.15

$$F_4(\text{AKD}, \text{O}; \text{Hmong}, \text{Han}) = 0 \quad (\text{S } 1.22)$$

By S 1.19, 1.21, and 1.22, we have

$$F_4(\text{Second\_MSEA}, \text{O}; \text{Hmong}, \text{Han}) \gg 0 \quad (\text{S } 1.23)$$

That is consistent with the result in Table 2. This completes the proof. ■

##### 1.4.2. Second_MSEA closer to Amis than Hmong

Using AKD instead of AAN/Insular_AN, and given the result in Table 2, we have

$$F_4(\text{AKD}, \text{O}; \text{Hmong}, \text{Amis}) \ll 0 \quad (\text{S } 1.24)$$

Given S 1.5 and S 1.20, we have

$$F_4(\text{Second\_MSEA}, \text{O}; \text{Hmong}, \text{Amis}) \ll 0 \quad (\text{S } 1.25)$$

This is consistent with the result in Table 2. This completes the proof. ■

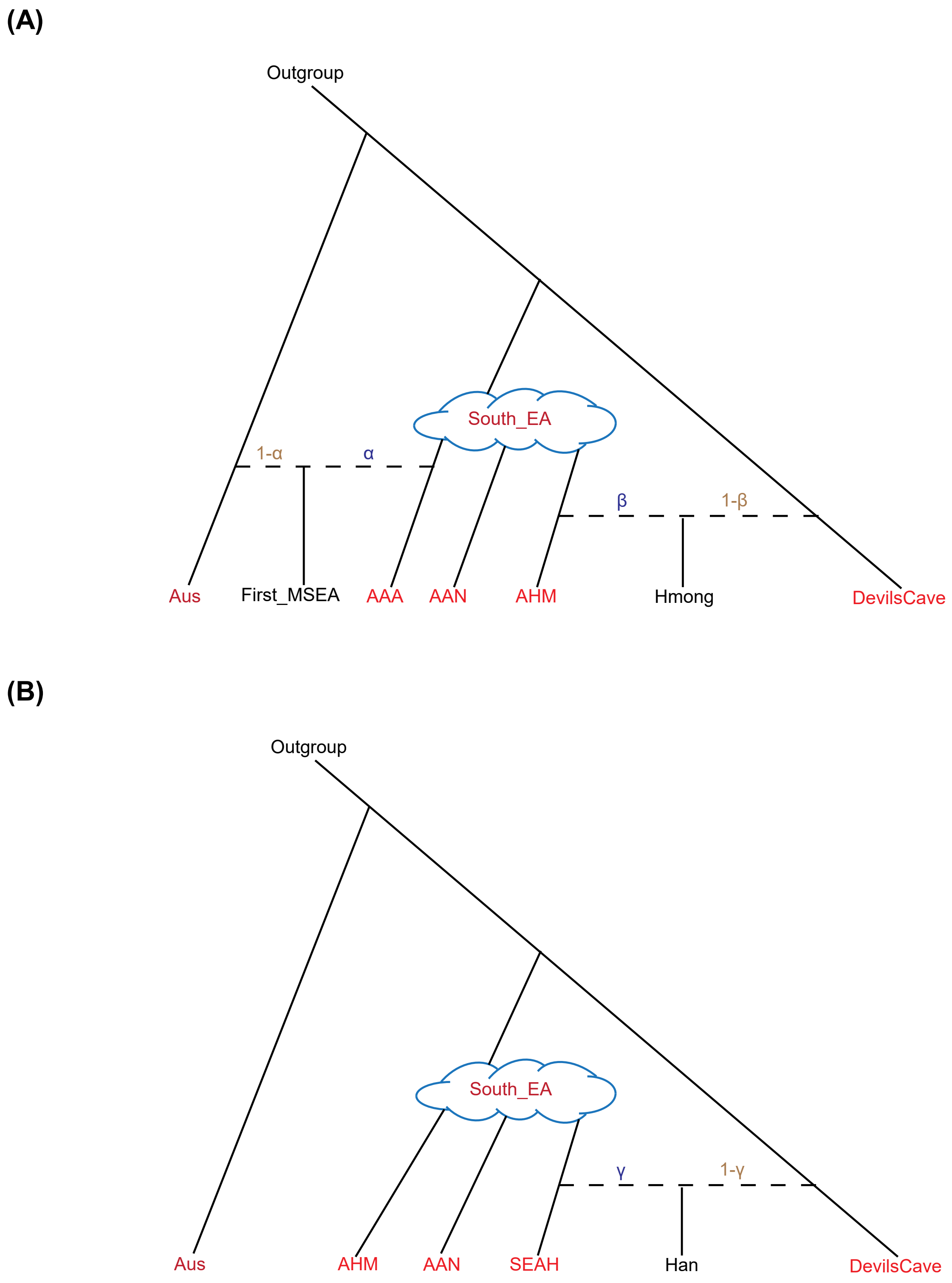


**SI Figure 1.1. | Sketch of the phylogenetic models.**
